# Posteromedial cortical networks encode visuomotor prediction errors

**DOI:** 10.1101/2022.08.16.504075

**Authors:** Ryosuke F. Takeuchi, Akinori Y. Sato, Kei N. Ito, Hiroshi Yokoyama, Reiji Miyata, Rumina Ueda, Konosuke Kitajima, Riki Kamaguchi, Toshiaki Suzuki, Keisuke Isobe, Naoki Honda, Fumitaka Osakada

## Abstract

Predicting future events based on internal models is essential for animal survival. Predictive coding postulates that errors between prediction and observation in lower-order areas update predictions in higher-order areas through the hierarchy. However, it is unclear how predictive coding is implemented in the hierarchy of the brain. Herein, we report the neural mechanism of the hierarchical processing and transmission of bottom-up prediction error signals in the mouse cortex. Ca^2+^ imaging and electrophysiological recording in virtual reality revealed responses to visuomotor mismatches in the retrosplenial, dorsal visual, and anterior cingulate cortex. These mismatch responses were attenuated when mismatches became predictable through experience. Optogenetic inhibition of bottom-up signals reduced a behavioral indicator for prediction errors. Moreover, cellular-level mismatch responses were modeled by Bayesian inference using a state-space model. This study demonstrates hierarchical circuit organization underlying prediction error propagation, advancing the understanding of predictive coding in sensory perception and learning in the brain.

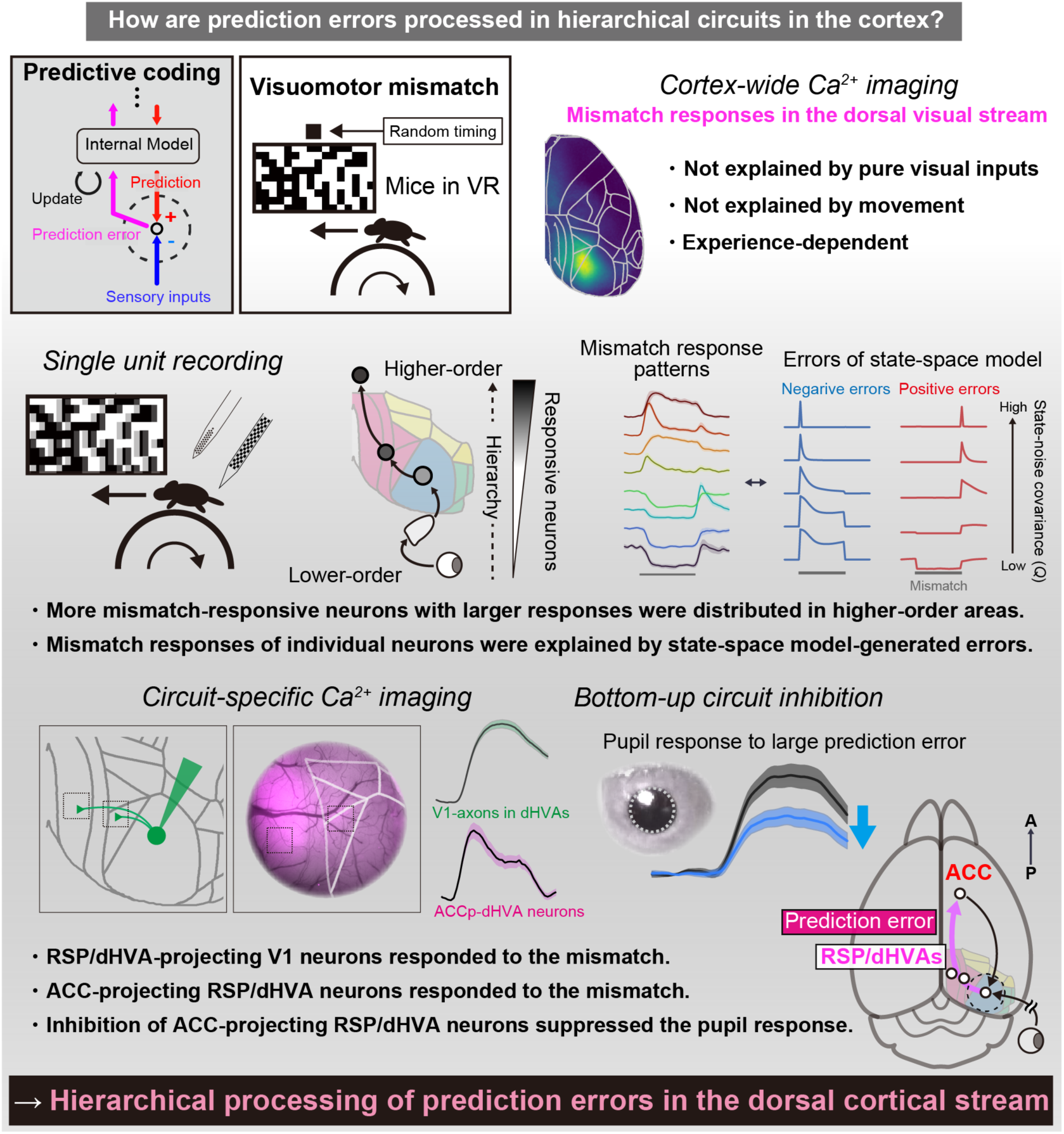

## Introduction

Animals can flexibly adapt to dynamic environments. The brain detects unexpected changes in sensory input, identifies them as prediction errors by comparing predicted sensory inputs to real-world observations, and updates an internal model of the environment^1–3^. According to the predictive coding hypothesis, top-down signals from higher-order brain areas convey sensory input predictions, whereas bottom-up signals from lower areas transmit prediction errors — the deviations between expected and observed sensory inputs^4,5^. In turn, the internal model encoded by higher-order circuits generates updated predictions to minimize future prediction errors. However, there is limited experimental evidence on how predictive coding is implemented within hierarchical neural circuits in the cerebral cortex.

Previous studies have demonstrated that top-down signals transmitted from movement-related areas are integrated with sensory inputs in lower-level sensory areas^3,6–10^. Among the higher-order areas in the brain, the anterior cingulate cortex (ACC) is a candidate for the areas that provide top-down, movement-related predictive signals to sensory areas^7,11^. The ACC is reciprocally connected with multiple sensory systems, such as higher-order visual and auditory cortex, and transforms sensory signals into goal-directed action signals to downstream areas^12–14^. Inactivation of the ACC impaired the error signal in the primary visual cortex (V1), suggesting that neurons in the V1 are capable of computing visuomotor prediction errors by utilizing top-down, movement-related signals provided by ACC^7^. However, little is known about how prediction error signals computed in lower-order areas propagate along the cortical hierarchy to the higher-order area and update the internal model. For instance, it is still uncertain how visuomotor prediction errors are processed at different levels of the cortical hierarchy and which regions outside V1 mediate prediction error propagation.

In this study, we aimed to determine how prediction error signals are hierarchically computed in the cerebral cortex, focusing on the visuomotor system. We utilized a virtual reality (VR) system to induce visuomotor prediction errors in animals by producing deviation between visual inputs and self-movement of animals^15^. Macroscopic Ca^2+^ imaging revealed that prediction error signals are processed by medial higher-order visual-related areas in retrosplenial (RSP), posteromedial (PM), and anteromedial (AM) visual cortex. Granger causality analysis results, prior neuroanatomical findings^16–18^, and theoretical considerations^4,5,19^ further suggested that prediction error signals are processed in hierarchical circuits along the dorso-dorsal cortical stream. Indeed, circuit-specific imaging and targeted optogenetic inhibition provided direct evidence for the flow of prediction error signals along the dorso-dorsal cortical stream. Prediction error responses in areas along this stream were hierarchical even at the cellular level, and the diverse neural response patterns were explained by a dynamic Bayesian inference computational model. Thus, we demonstrate that hierarchical circuits along the dorsal cortical stream process prediction errors, thereby revealing the neural implementation of predictive coding in the cerebral cortex.

## Results

### Regional encoding of prediction error signals in the medial cortex

To study prediction error signals in the cerebral cortex, we labeled neurons across cortical areas with the genetically encoded Ca^2+^ indicator jGCaMP7f and performed wide-field Ca^2+^ imaging in head-fixed mice running on a treadmill under a VR environment. An intravenous injection of a blood–brain barrier-permeable adeno-associated virus vector (AAV-PHP.eB) expressing jGCaMP7f under control of the neuron-specific human synapsin promoter into the mouse retro-orbital sinus introduced jGCaMP7f into neurons throughout the brain, including cortex, striatum, hippocampus, and cerebellum (Fig. 1A, B, Fig. S1). Histological analysis showed that only a small proportion of the jGCaMP7f-expressing cells in V1 and PM were immunopositive for the inhibitory neuron subtype markers parvalbumin (5.9% in V1, 8.1% in PM), somatostatin (1.1% in V1, 0.8% in PM), or vasoactive intestinal peptide (0.0% in V1, 0.5% in PM), indicating biased labeling of excitatory neurons, consistent with the previous report^20^.

**Figure 1.**
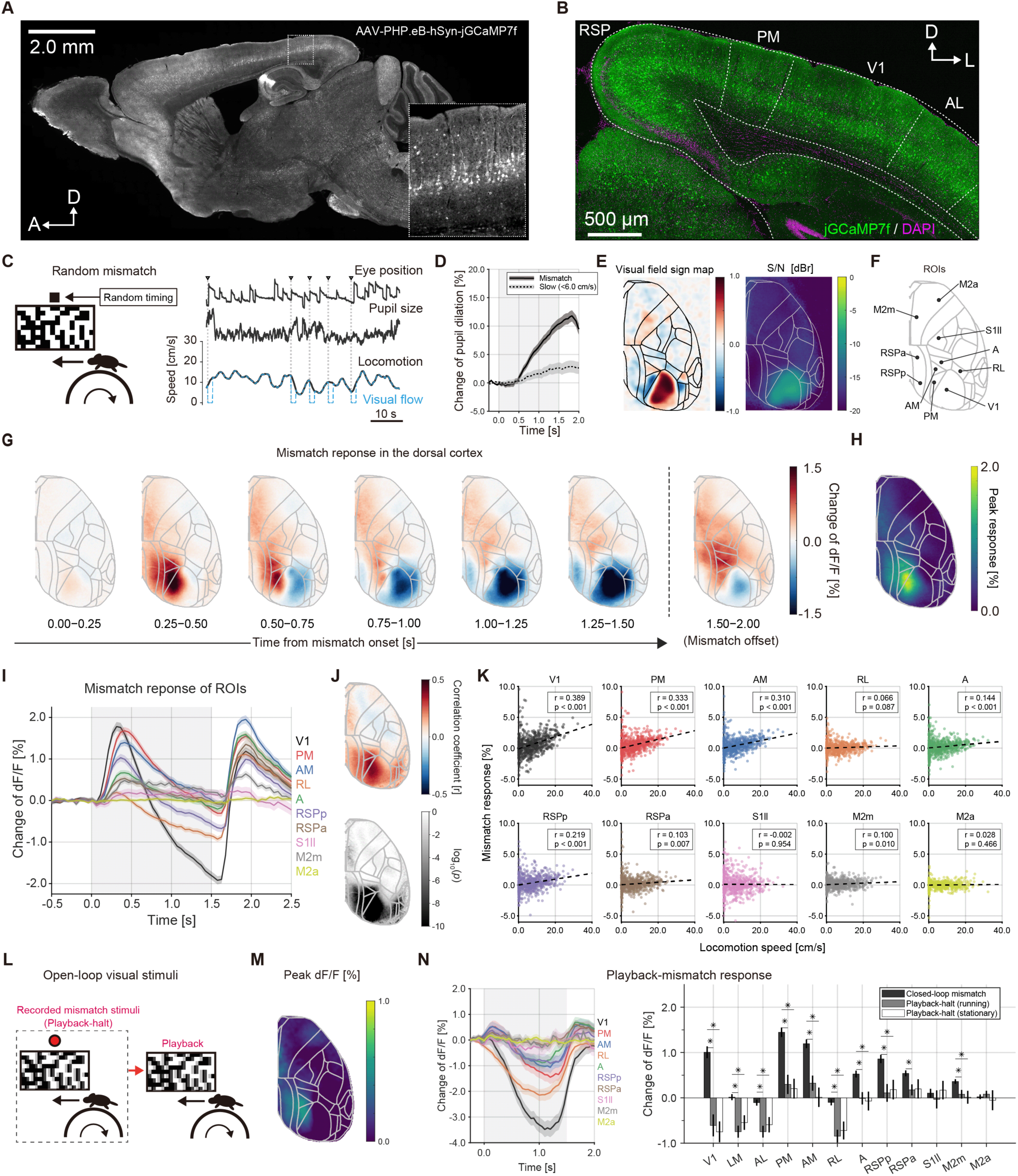
Visuomotor mismatch responses in dorsal cortical areas. (A). Sagittal section of mouse brain showing widespread cortical and subcortical expression of the genetically encoded Ca^2+^ indicator jGCaMP7f in the AAV-PHP.eB-infected brain. Inset: Magnified view of jGCaMP7f-expressing neurons in primary visual cortex (V1). Arrows indicate anterior (A) and dorsal (D) axes. (B). Coronal section from the jGCaMP7f-expressing brain. Arrows indicate dorsal (D) and lateral (L) axes. (C). Left: Schematic of the virtual reality (VR)-based visuomotor mismatch paradigm. Right: Example time series of mismatch and extracted behavioral parameters. Arrowheads indicate the timing of mismatch onset. (D). Time course of the pupil dilation response: Solid line represents the trial-averaged response to mismatch events, and dashed line illustrates the trial-averaged response when mice exhibited minimal locomotion (<6.0 cm/s) at the mismatch onset. Shaded areas: SEM. (E). Left: Averaged visual field sign map. Right: A signal-to-noise ratio (S/N) map (n = 6 mice) during visual stimulation with a moving and flickering checkerboard pattern. (F). Locations of 10 region of interest (ROIs) on the top view of the ACCF atlas. (G). The trial-averaged response to visuomotor mismatch events (326 trials from nine mice). (H). Response magnitude map during the mismatch period (0.0−1.5 s). (I). Averaged response traces from the 10 identified cortical areas. Shaded region indicates the mismatch periods (0.0−1.5 s). Line traces represent means ± standard error of the mean (SEM). (J). Top: Pseudocolor map of Pearson’s correlation coefficient between locomotion speed at the mismatch onset (mean of −0.25−0.25 s) and the neural response magnitude at all pixels in the ACCF. Bottom: Grayscale map of corresponding *p*-values. (K). Scatter plot of locomotion speed and mismatch responses in the 10 cortical areas. (L). Schematic of open-loop stimulus presentation. (M).Peak response map to playback-mismatch events (n = 4 mice, locomotion speed > 6.0 cm/s). (N). Left: Averaged response traces from the 10 cortical ROIs. The shaded region indicates the mismatch period (0.0−1.5 s). Line traces represent means ± SEM. Right: Comparison of responses to closed-loop mismatch, playback-halt during locomotion, and stationary state conditions. Asterisks show statistical significance in each area (*p* < 0.01, by Mann–Whitney U test).

The VR system allows for precise manipulation of the relationship between locomotion and resulting visual feedback for selective delivery of sensorimotor mismatches (prediction errors) after habituation of mice in VR (Fig. 1C). To determine how mice respond to visuomotor prediction errors, we presented unexpected visual feedback perturbations of the closed-loop relationship between visual flow and locomotion by randomly stopping the visual flow for 1.5 s while mice running on the treadmill (referred to as mismatches)^15^. During the mismatch event, pupil dilation occurred reliably in response to these mismatches (Fig. 1C-D), consistent with previous studies showing that pupil dilation responses reflect arousal or surprise due to visuomotor mismatches^15,21,22^. To filter the calcium-independent signal from neural activity, we performed wide-field Ca^2+^ imaging with a fluorescence normalization scheme on the basis of wavelength multiplexing (Fig. S2, See methods)^23,24^. We computed a visual field sign map, resting-state connectivity map, and principal component analysis map as references for registering the brain activity map of individual animals to that in the Allen Common Coordinate Frameworks (ACCF, Fig. 1E and S3, Supplementary Videos 1 and 2)^25^. We defined 10 regions of interest (ROIs) based on the ACCF (Fig. 1F). Of these, RSP was subdivided into anterior and posterior portions (RSPa and RSPp) based on anatomical differences in neural circuit inputs and outputs, as determined by tracer injection (Fig. S4)^26^.

Wide-field Ca^2+^ imaging revealed that visuomotor mismatches evoked widespread and sequential cortical activities (Fig. 1G, Supplementary Video 3). Distinct changes in the jGCaMP7f signal were detected in medial visual areas, including V1, PM, AM, and RSPp. In contrast, the rostrolateral (RL) and lateral visual areas did not exhibit distinct positive peak Ca^2+^ responses (Fig. 1H, I). As a negative control, fluorescent signals of green fluorescent protein (GFP)-expressing neurons were also measured, and no such changes in fluorescence during mismatch were detected (Fig. S5). We then investigated whether these signals encode the magnitude of prediction errors by calculating the correlation between mismatch response magnitude and prediction error magnitude^15^ . For this analysis, the magnitude of the prediction error was measured as the locomotion speed at mismatch onset (mean speed from −250 to 250 ms, threshold = 0.02 cm/s). The mismatch response magnitudes in medial higher-order visual areas, such as PM and AM, were significantly correlated with locomotion speed (Fig. 1J, K). Notably, the RSPp also exhibited a larger correlation between mismatch response magnitude and locomotion speed than RSPa (Fig. 1K, RSPp: *r* = 0.219, RSPa: *r* = 0.103). In contrast, such correlations were not observed in parietal association areas, such as the RL and anterior visual area. Furthermore, correlations with movement-related variables during the mismatch event, such as changes in locomotor speed and facial motion^24,27^, were much weaker and could not account for the variability in mismatch responses compared to locomotor speed at mismatch onset (Fig. S6). From these results, we conclude that medial higher-order areas, such as PM, AM, and RSPp, encode the occurrence and magnitude of visuomotor prediction errors. The mismatch event not only includes a prediction error (difference between expected and actual sensory input based on locomotion) but also a visual flow speed change. Prior studies have highlighted both the slower speed preference of visual flow in PM and the amplification effect of locomotion on sensory responses in visual areas^9,28,29^. In addition, we observed that dorsal cortex activity increased at the onset of running, in the absence of visual feedback (Fig. S7). Therefore, the mismatch responses may be explained by amplification of responses to the visual flow velocity change due to animal locomotion rather than the prediction error *per se*^9,30^. To rule out this possibility, we conducted wide-field Ca^2+^ imaging on mice under both the closed-loop mismatch condition and an open-loop condition in which mice were presented with playback-mismatch visual stimuli that were not coupled to locomotion speed (Fig. 1L). Consistent with previous reports in V1 neurons^15^, all mismatch-responding areas showed smaller response magnitudes to playback-halt (playback of mismatch event) stimuli than mismatch stimuli, regardless of the locomotion speed (Fig. 1L–N). The difference in playback responses between running and stationary conditions was relatively small (*p* > 0.05 in all ROIs). Thus, it is unlikely that the visual flow alone or modulation by locomotion accounts for the mismatch response under the closed-loop condition.

### Neural and behavioral mismatch responses are experience-dependent

To examine whether pupil and neural mismatch responses are modulated by prediction formed by prior experience, we compared neural responses to “predictable” and “unpredictable” visuomotor mismatches while maintaining identical locomotion speeds. If neural activity associated with the mismatch response reflects prediction error, then this response should be attenuated when the mismatch event is highly predictable. To test this notion, we established a semi-closed-loop VR (SCL-VR) condition in which the perturbation of visual flow feedback occurred only when the locomotion speed exceeded a threshold of 12.0 cm/s, thereby triggering a self-induced mismatch (Fig. 2A). In contrast to the random timing of mismatches in our initial experiments (Fig. 1), the timing of these self-induced mismatches is predictable for a mismatch-experienced (ME) group trained in the SCL-VR. However, for the normal VR-experienced (NE) group trained in the regular closed-loop VR (the control group in this experiment), the timing of the self-induced mismatch in the SCL-VR is unpredictable (Fig. 2B). In both groups, the visuomotor mismatch responses were mainly attributed to predictions based on prior experience during five habituation sessions, and consequently the mismatch response magnitude was expected to be smaller in the ME group than the NE group (Fig. 2C).

**Figure 2.**
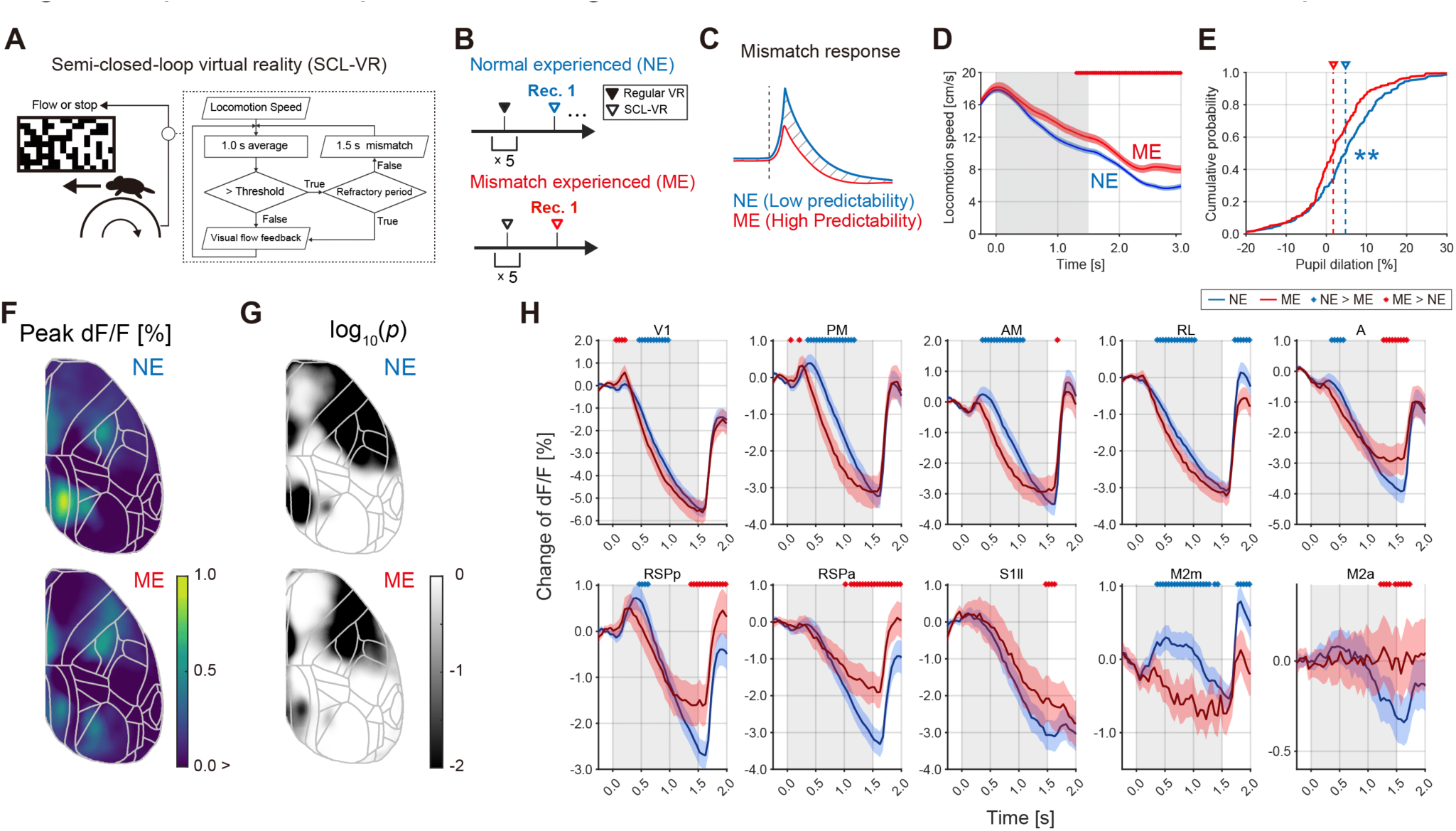
Experience-dependent changes in self-induced visuomotor mismatch responses. (A). Schematic of semi-closed-loop virtual reality (SCL-VR), with inset showing the logic flow of visual feedback. (B). Habituation history of two groups of mice differing in mismatch experience, the normal experienced (NE) group and mismatch experienced (ME) group. (C). Hypothetical neural responses to mismatch events by NE and ME mice. (D). Locomotion speed of mice during self-induced mismatch events. Shaded region indicates the mismatch period (0.0−1.5 s). Line traces represent means ± SEM. Dots show time bins with statistical significance (*p* < 0.01, by one-tailed unpaired *t*-test). (E). Distribution of pupil response magnitudes on each trial (*p* < 0.01, by Mann–Whitney U test). (F). Response peak magnitude maps during the mismatch period (0.0−1.5 s). (G). Statistical significance of self-induced mismatch response between NE and ME group mice (average of 0.0–0.5 s vs. −0.5–0.0 s, *p* < 0.05, by one-tailed Signed–Rank test). (H). Trial-averaged response traces from 10 ROIs. Shaded region indicates the mismatch period (0.0−1.5 s). Blue plots show the data from NE mice and red plots the data from ME mice. Each plot is shown as a mean ± 95% confidence interval. Dots indicate time bins with significant difference (*p* < 0.05, by bootstrap test). Dot color indicates the alternative conditions (blue: NE > ME; red: ME > NE).

During recording sessions, the average locomotion speed in the early phase of the mismatch period (0.0 to 1.2 s) did not differ significantly between NE and ME groups (Fig. 2D). In the late phase of the mismatch periods (>1.2 s), however, the NE group exhibited significantly slower locomotion speed and larger pupil dilation than the ME group (Fig. 2D, E and S8A). These results suggest that the ME group acquired prior information based on experience in the SCL-VR, resulting in reduced prediction errors to self-induced mismatch events compared to the inexperienced NE group.

To examine if these effects are associated with neural response magnitude in specific cortical regions, we compared the neural responses to the self-induced mismatches between the NE and ME groups using wide-field Ca^2+^ imaging (Fig. 2F). While positive mismatch responses were observed in both groups across wide regions of dorsal cortex, such as medial secondary motor cortex (M2m), RSPp, AM, and PM (Fig. 2F, G), response magnitudes were uniformly significantly smaller in the ME group during the early period of the mismatch (0.0 to 0.75 s) (Fig. 2H, S8B and C). To further examine the effect of experience on mismatch responses, we performed additional recordings in NE mice following each of four additional training sessions in the SCL-VR. In the recording second session, NE mice showed reduced behavioral responses in locomotion speed and pupil dilation compared to the first session (Fig. S9A–C) as well as reduced neural activity in the V1, PM, AM, and RSPp (Fig. S9), indicating that mismatch responses observed in the medial higher-order cortical areas are contingent on the predictability of mismatch events.

### Mismatch responses of individual neurons are explained by the visuomotor prediction model

Our wide-field Ca^2+^ imaging results (Fig. 1 and 2) and previous anatomical and physiological findings on the cortical visual system^16–18,31^ led us to reason that visuomotor prediction errors are hierarchically processed in a dorsal cortical pathway (Fig. 3A and B). To assess regional differences in mismatch responses at the single-neuron level, we performed extracellular recordings from cortical areas exhibiting robust mismatch responses by wide-field imaging (V1, PM, RSPp, and ACC, including 338 single units in total from n = 9 mice, Fig. 3A, B and Fig. S10) as well as from potentially relevant subcortical areas dorsal lateral geniculate nucleus (dLGN) and dorsal or ventral hippocampal region (dHIP, vHIP), 501 single units in total from n = 4 mice, Fig. S10). These recordings were performed from mice in two or three types of VR tracks after the session for area identification with intrinsic signal optical imaging^32,33^ (Fig. 3C). Track 1 presented a corridor with landmarks, Track 2 a corridor without landmarks, and Track 3 a dark environment. Remarkably, recorded neurons showed highly stable mismatch responses across trials, and there were no significant magnitude differences in V1, PM, RSPp, and ACC between Track 1 and Track2 conditions (all *p* > 0.05 by two-sided Sign-rank test), indicating that the mismatch responses are not largely affected by spatial prediction. Next, we examined whether mismatch response patterns change along the dorsal visual pathway. In the dLGN, the earliest visual processing stage in the brain, 15.5% of units showed a significant positive response to mismatch; in V1, 30.0 % of units showed a significant positive response to mismatch (Fig. 3I). Further, the proportion of responsive neurons gradually increased along the cortical hierarchy, as mean responses were greatest in PM and RSPp, lowest for dLGN, and of intermediate magnitude in V1 and ACC (Fig. 3J). The response magnitudes in V1, PM, and RSPp were also significantly larger than that in dLGN (*p* < 0.01, Dunnett’s test). These results suggest that areas at the higher level of the hierarchy may be more engaged in predictive processing.

**Figure 3.**
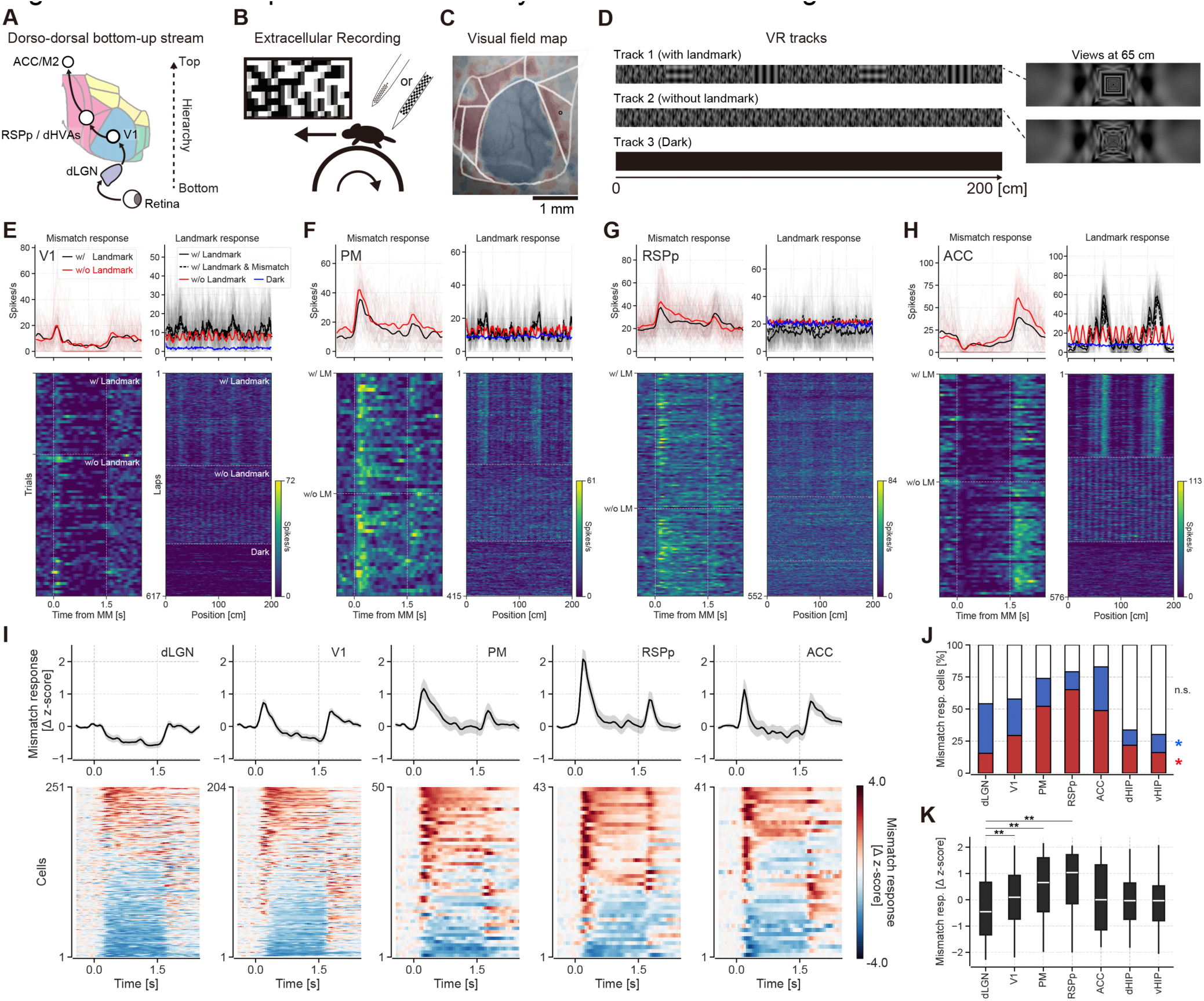
Mismatch responses measured by extracellular recording. (A). Hypothetical visuomotor error processing along the hierarchical structure in the dorsal cortex. (B). Schematic illustration of single-unit electrophysiological recording from mice under VR. (C). Visual field map for identification of recording sites. (D). VR tracks presented during recordings. (E). Top left: Trial-averaged mismatch response of a representative V1 neuron. Bottom left: Mismatch response on each trial, sorted by VR track and locomotion speed. Top right: Firing rate traces across positions in the virtual track. Bottom right: Lap-by-lap firing rate. The dashed line in the matrix indicates the switching of VR track (Track 1, Track 2, Track 3, respectively). (F). Same as in (E) for a representative PM neuron. (G). Same as in (E) for a representative RSP neuron. (H). Same as in (E) for a representative ACC neuron. (I). Mismatch responses of neurons in the dLGN, V1, PM, RSPp, and ACC (in Track 1 and Track 2). Top: Population average of mismatch response. Shaded region indicates the SEM. Bottom: Mismatch response of individual neurons in each area. Neurons were sorted by mismatch response magnitude. (J). Bar charts of mismatch responsive neurons in the dLGN, V1, PM, RSPp, ACC, dorsal hippocampus (dHIP), and ventral hippocampus (vHIP). Cells were classified according to one-sided sign rank test (*p* < 0.05). The red area indicates positive responsive cells. The blue area indicates negative responsive cells. (K). Mismatch response magnitudes across the dLGN, V1, PM, RSPp, ACC, dHIP, and vHIP. Each dot indicates an individual neuron. Asterisks indicate statistically significant differences (** *p* < 0.01, vs dLGN, by Dunnett’s test).

Notably, the pattern of neuronal mismatch responses was more diverse than presented in previous studies^15,34^. We next classified the mismatch response patterns of individual units using an unsupervised learning scheme (time series K-means clustering). This analysis revealed eight distinct clusters (Fig. 4A and B). Moreover, mismatch responses of individual neurons were not solely limited to simple increases and decreases in neural activity. This response complexity suggests that the neural circuits encoding of prediction errors do not merely perform simple arithmetic operations.

**Figure 4.**
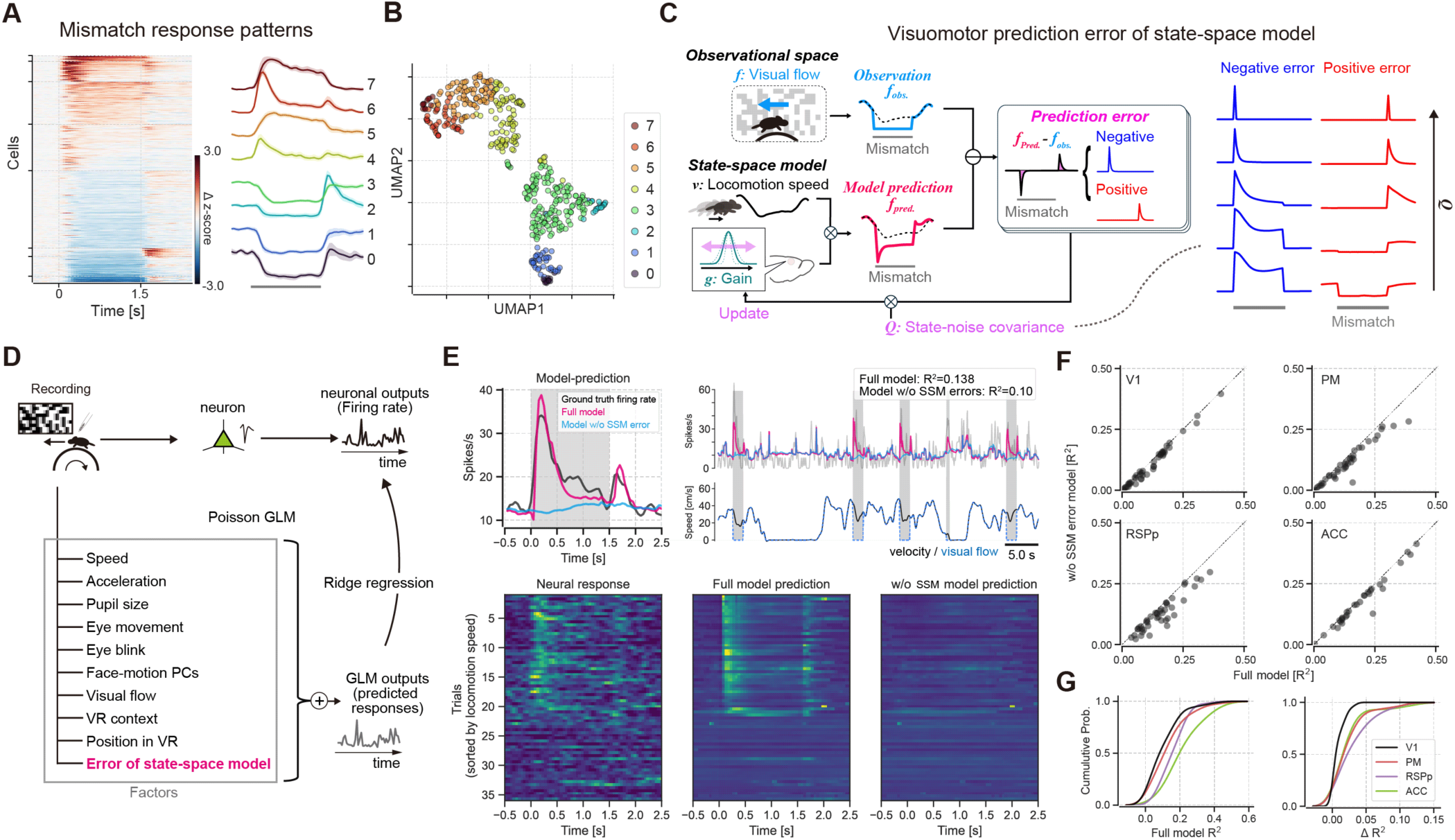
Errors by the visuomotor prediction model explain neural mismatch responses. (A). Variable mismatch response patterns in the dLGN, V1, PM, RSP, ACC, dHIP, and vHIP. Left: Response patterns of mismatch-responsive cells (sorted by time-series K-means clustering). Right: Mean mismatch response of each cluster. Shaded areas indicate SEM. Gray bar indicates the mismatch periods. (B). UMAP projection embedding the mismatch response pattern of individual neurons. Cluster labels (0-7) correspond to those in (A). (C). Schematic of the visuomotor predictive state-space model (SSM). Gray bar indicates the mismatch period. (D). Schematic of Poisson generalized linear model (GLM) analysis. (E). Model fitting results for an example PM neuron. Top left: Averaged mismatch response of the PM neuron (gray line), Full model prediction (blue line), and prediction of the model without visuomotor prediction errors calculated by the SSM (red line). Gray shaded region indicates the mismatch periods. Top Right: example time course of actual firing and fitting result. Bottom: Mismatch response pattern matrix of neural responses and the predictions of each model. (F). Comparison of fits for the full model and the partial model without SSM variables. Dots indicate individual cells in V1, PM, RSPp, and ACC. (G). Left: GLM fit quality, measured as the R-squared value (R^2^). Right: Distribution of ΔR^2^ (R^2^ _Full model_ - R^2^ _w/o SSM model_) for each recording site.

To test further examine if response patterns of individual neurons are consistent with predictive coding scheme, we compared neural mismatch response to visuomotor prediction errors estimated by a state-space model (SSM)^1^ that predicts visual flow speed from locomotion speed and visual flow histories of mice (Fig. 4C). During closed-loop periods, this SSM model accurately predicted the incoming visual flow speed, while during the mismatch periods, the model detected the visuomotor prediction errors and updated its state accordingly. Visual flow speed prediction in the SSM involves estimating the latent state variable *gt* parameterized by the state noise covariance (*Q*). In this model, the value *Q* acts as a hyperparameter that determines the sensitivity of the prediction error in the SSM, with larger values indicating that the model can quickly adapt to large variations in the state transition (Fig. 4C, see Methods for details). This model successfully computed the visuomotor prediction error as the mismatch between the observed and the expected visual flow based on the locomotion speed during the recording session (Fig. S11). The error between predicted and actual visual flows varied depending on *Q*; that is, various patterns of visuomotor prediction errors in SSM can be produced by this hyperparameter. This finding further implies that the variable Q in SSM can be considered a control parameter that determines how much the state is updated using the received information. Notably, the model generated not only simple positive and negative response patterns, but also transient and persistent response patterns to mismatches depending on different hyperparameters (Fig. 4C). Furthermore, these model-generated patterns recapitulated the distinct spiking response patterns in neural activity of individual neurons to mismatch (Fig. 4A and B).

To test whether the visuomotor prediction errors estimated by the SSM can explain neuronal responses, we employed a generalized linear model (GLM) designed to evaluate the contribution of SSM-generated prediction error information to actual neural responses of individual cells. The GLM used variables associated with mouse behaviors and sensory inputs such as visual flow and landmarks in the VR, in addition to the prediction error information generated by the SSM (Fig. 4D). To assess relative contribution of each behavioral variable to the GLM performance for individual neurons in V1, PM, RSPp, and ACC, we evaluated the effects of omitting a single feature (SSM-generated errors) from the full model and evaluated the GLM performance by comparing the partial model with the full model (Fig. 4E–G). The omission of SSM-generated prediction errors decreased GLM performance across all four brain regions (Fig. 4 F and G), indicating that prediction errors are crucial for explaining neural activities in V1 and higher-order areas PM, RSPp, and ACC. This finding suggests that the neural mismatch response does not merely represent sensory inputs, self-movements, or the difference between the efference copy and the observation, but rather the difference between prediction and observation, with the prediction being dynamically updated based on past errors. Notably, while the contribution of SSM errors to neuronal responses was significant in V1, it was even more pronounced in higher-order areas (PM, RSPp, and ACC, Fig. 4G), supporting our hypothesis that prediction errors are hierarchically processed along the cortical hierarchy.

### Hierarchical propagation of bottom-up error signals along the dorso-dorsal visual stream

We then investigated the transmission route of error signals among across dorsal cortical areas by applying correlation analysis between cortical areas where neural activity was simultaneously measured with wide-field Ca^2+^ imaging. There were significant differences in large-scale correlations between cortical areas under conditions with distinct behavioral states and sensory inputs (Fig. 5A). While the overall magnitudes of the correlations decreased when mice were running in the VR environment and the pattern was roughly preserved between during the closed-loop and mismatch periods, individual areal correlation coefficients increased across visual cortical areas during mismatches (Fig. 5A, B). To quantify the direction and degree of information flow between areas, we computed Granger causality during mismatch and closed-loop periods^35,36^. Despite dynamic changes in visual flow speed during closed-loop periods, Granger causality from V1 to PM remained lower than during mismatch periods (Fig. 5C and Fig. S12A). We also applied Granger causality analysis to the self-induced mismatch responses of NE and ME group mice in the SCL-VR under similar locomotion speeds and visual flow feedback conditions (Fig. 2) and found reduced Granger causality from V1 to higher-order visual areas in ME mice (Fig. S12B). These results suggest the existence of bottom-up error signal flow from V1 to higher areas during the mismatch periods.

**Figure 5.**
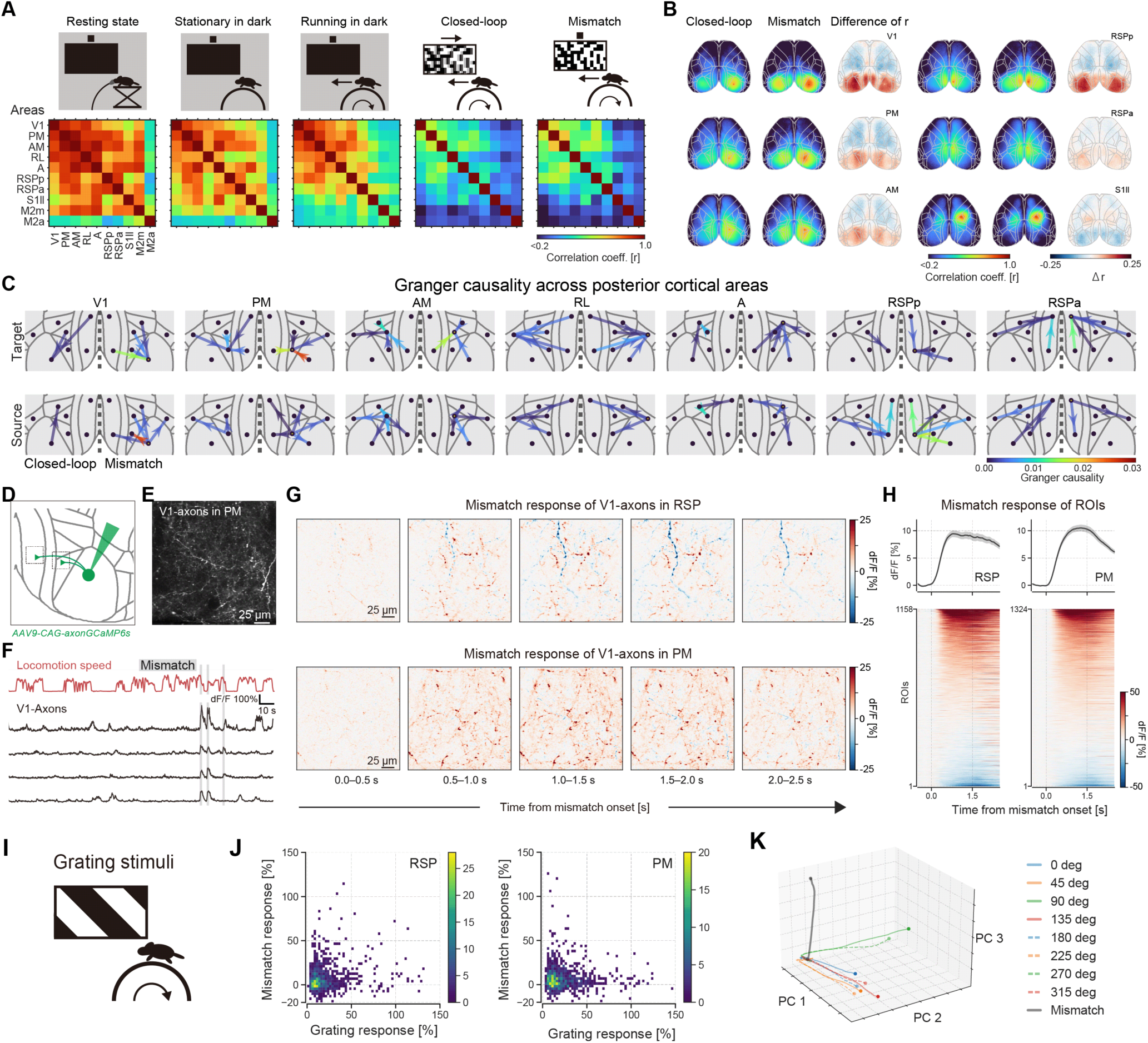
Error signal flow across dorsal cortical areas. (A). Schematic of five distinct behavioral and environmental contexts. Corresponding matrices show pairwise linear correlation coefficients of activity among the 10 defined ROIs. (B). Seed-based correlation maps of V1, PM, AM, RSPp, RSPa, and S1ll activity in the closed-loop and mismatch periods, and the differences in correlation coefficient between conditions. (C). Granger causality between posterior cortical areas during closed-loop and mismatch periods. Closed-loop results are visualized in the left hemisphere and mismatch results in the right hemisphere. Statistically significant flow edges are shown (*p* < 0.01). Open circle indicates seed-ROI. (D). Schematic of labeling and imaging of V1 axons. (E). Example *in vivo* two-photon image of GCaMP6s-labeled V1 axons in PM (F). Example time course of locomotion speed, mismatch events and dF/F of ROIs in PM. (G). Pixel-based trial-averaged mismatch responses of V1 axonal boutons in example recording sites in RSP and PM (H). Trial-averaged mismatch responses from ROIs in RSP and PM. ROIs were sorted by mismatch response magnitudes. (I). Schematic of visual stimulation (drifting rectangular grating). (J). Relationship between grating response (preferred direction) and mismatch response amplitude visualized by density plot. Colors indicate the counts of ROIs in each bin. (K). Projection of the population responses to visual stimuli and mismatch events into 3-dimensional PCA space. Trajectories indicate population dynamics during stimulus presentation (0.0-1.5 s). Circles indicate offset of stimulus.

To directly examine this functional connectivity at the circuit level, we performed two-photon Ca^2+^ imaging of axonal boutons from V1 neurons in the RSP and PM following antero-grade labeling with axon-GCaMP6s (Fig. 5D and E). Injection sites and recording sites were identified by intrinsic signal optical imaging and post-hoc anatomical validation. Figure 5F–G shows examples of mismatch response time courses. The Ca^2+^ signal time courses of representative boutons clearly revealed positive and negative responses to mismatch events (Fig. 5G). ROIs were then defined to quantify mismatch response of axons in RSP and PM (n = 1158 and 1324 ROIs from n = 3 and 3 mice, respectively, Fig. 5H). A significant proportion of axonal boutons in PM and RSP responded to mismatch events (Fig. 5H; 34.5% in RSP, 43.7% in PM, *p* < 0.05). To investigate the relationship between mismatch responses and visual response properties, we additionally measured the visual responses to drifting grating stimuli in a subset of recording sessions (Fig. 5I, n = 1174, 1121 ROIs, respectively). Notably, the distribution of neuronal response properties projected onto grating and mismatch response axes were nearly orthogonal (Fig. 5J). Furthermore, we examined whether the population neuronal dynamics for mismatch responses were separable from those for visual responses to various grating stimuli, using a principal component analysis (PCA) to decompose the high-dimensional neural dynamics, and demonstrated that the neuronal population dynamics of mismatch responses were clearly separable from those of visual responses to grating stimuli (Fig. 5K). These axon imaging analyses, together with the macroscopic level results, indicate that these error-responsive V1 neurons transmit prediction error signals to higher-order areas, and are distinguishable from neurons sending bottom-up visual information.

Finally, we examined the feedforward propagation of visuomotor prediction error signals from PM and RSP to motor-related areas, as such signals are considered necessary for updating the internal model and refining future predictions^4^. For this purpose, we retrogradely labeled ACC-projecting neurons with AAV2retro-tdTomato in Thy-1-GCaMP6f transgenic mice^37^ and then performed two-photon Ca^2+^ imaging from the RSP and medial higher visual areas (PM and AM). Recording sites were determined by retinotopic mapping (Fig. 6B)^32^. A considerable number of ACC-projecting RSP and PM/AM neurons (tdTomato^+^) in layer 2/3 exhibited a significant positive response (24.2% and 31.1%, respectively) to mismatch events (Fig. 6D-F). Further, the mismatch response magnitude of the tdTomato^+^ neurons was positively correlated with locomotion speed (Fig. 6G), and the spatial tuning profile and playback-halt response of these neurons did not account for the mismatch response magnitudes (Fig. S13 and S14). These results indicate that ACC-projection neurons in RSP and PM/AM directly transmit the prediction error signals to ACC.

**Figure 6.**
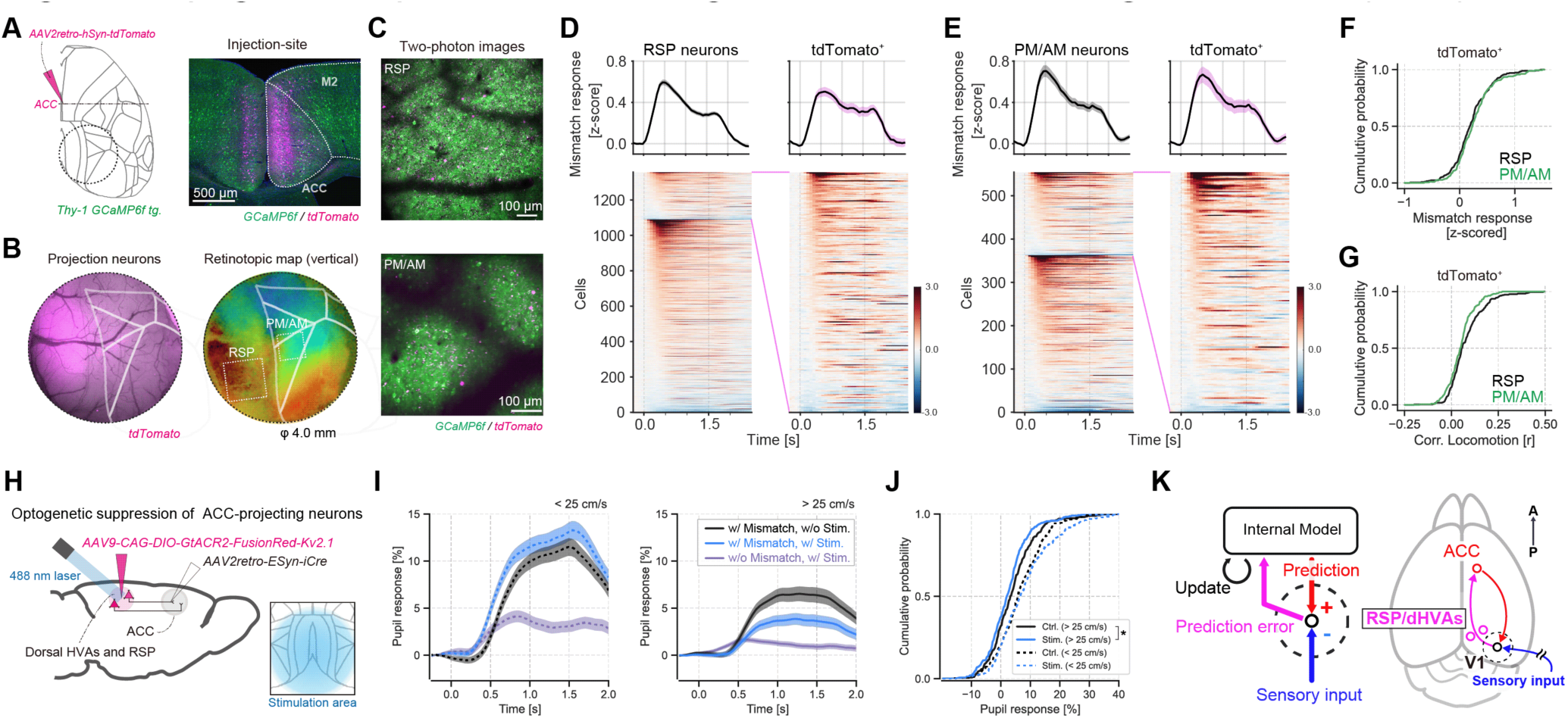
Propagation of prediction error signals to the anterior cingulate cortex (ACC) (A). Labeling of ACC-projecting neurons by AAV into the ACC of Thy1-GCaMP6f transgenic mice (left) and a confocal image of the injection site (right). The circle represents a typical glass window position for *in vivo* two-photon imaging. (B). Fluorescent images of recording sites (left) overlaid with the retinotopic mapping result (right). The real boundary of the dorsal visual areas is outlined. Boxes indicate the example imaging site in (A). (C). *In vivo* two-photon images of the RSP and PM/AM. (D). Trial-averaged mismatch response from the RSP. Left: Averaged response of all recorded neurons in the RSP (n = 1092 tdTomato^-^ cells, n = 265 tdTomato^+^ cells from 4 mice). Right: Averaged response in ACC-projecting RSP neurons. (E). As in (D) for PM/AM neurons (n = 361 tdTomato^-^ cells, n = 193 tdTomato^+^ cells from 4 mice). (F). Mismatch response magnitude (mean of 0.0-1.5 s) distribution of ACC-projecting neurons. (G). Correlation coefficient between locomotion speed and neural activity of ACC-projecting neurons under the darkness. (H). Optogenetic silencing of ACC-projection feedforward circuits. Inset shows laser-illuminated area. (I). Time course of pupil dilation response to mismatch events for relatively slower trials (left) and faster trials (right). Shaded regions indicate the SEM. (J). Distribution of pupil response magnitude (Ctrl. vs. Stim.: *p* = 0.227 in <25cm/s trials; *p* = 0.010 in >25cm/s trials, by Mann–Whitney U-test). (K). Schematic of the hierarchical predictive coding process of visuomotor signals.

Next, to investigate whether these bottom-up circuits mediate the perception of prediction error, we performed loss-of-function analysis on pathways from posteromedial cortical areas, including the PM, AM, and RSP, to the ACC by optogenetic inactivation. We expressed the light-gated chloride channel GtACR2 in ACC-projecting neurons of the PM/AM and RSP by injecting retrograde Cre-expressing AAV (AAV2retro-hSyn-iCre) into the ACC and Cre-dependent AAV expressing GtACR2 (AAV9-CAG-DIO-GtACR2-FusionRed-Kv2.1) into the PM/AM and RSP (Fig. 6H and S15). Optogenetic inactivation of the ACC-projecting neurons in posteromedial cortical areas (PM, AM, and RSP) during mismatch periods significantly reduced the pupil dilation response to mismatch events in trials with large prediction errors (Fig. 6I, J). As a control, optical stimulation alone without visuomotor mismatch elicited only a slight and insignificant pupil dilation (Fig. 6I). These effects were consistent across individual animals (Fig. S16). Taken together, our results indicate that dorsal posterior-frontal cortical circuits propagate visuomotor prediction errors, supporting the theory of hierarchical predictive coding in the brain (Fig. 6K).

## Discussion

In this study, we elucidated the functional dynamics and neural circuit structure of prediction error encoding and propagation from lower-to higher-order areas through a cortical hierarchy. Multiscale recording and neural manipulation approaches combined with the visuomotor mismatch paradigm using VR revealed hierarchical encoding of experience-dependent visuomotor error signals along the dorso-dorsal visual pathway to the frontal cortex. Prediction error responses at the cellular level varied markedly among neurons and their statistical modeling further suggests that the mismatch response diversity results from prediction error computations according to complex probabilistic neural processing rather than simple arithmetic comparisons between sensory and internal model prediction signals. Inhibition of bottom-up signals from the medial higher-order regions RSP, PM, and AM to the ACC reduced the mismatch-induced pupil dilation response, a possible behavioral indicator of large prediction errors, suggesting that bottom-up signals within the cortical hierarchy are required for error detection. We conclude that the hierarchical processing of prediction error signals from V1 through medial higher-order posterior areas to the frontal motor area, a primary source of predictions, is critical for updating internal models and generating future predictions in the brain.

### Hierarchical propagation of bottom-up prediction error signals

Our results are consistent with the principles of both hierarchical predictive coding and the well-established hierarchical properties of visual processing^16,18,38–40^. Axonal imaging demonstrated that the population representations of basic visual inputs and prediction error information are nearly orthogonal in low-dimensional space, suggesting that the neurons conveying these two types of information are segregated, allowing downstream neurons to distinguish between visual feature information and prediction error signals (Fig.5). Future investigations are warranted to link mismatch-responsive neurons (function) to molecularly defined cell types (molecular profiles) and projection targets (connectivity) to understand circuit interactions of functionally distinct pathways^34,41,42^. Further, elucidating the interactions among parallel and hierarchical pathways is essential for a general understanding of predictive processing in the brain.

Detecting unexpected visual input during movement is crucial for visually guided motor control, perception, navigation, and learning. Studies in primates and rodents have indicated that visuomotor error signals in motor cortices play a pivotal role in driving adaptive motor learning^43,44^. Clinical and disease model studies have shown that impairments in medial higher cortical areas, including the dorsal higher-order visual areas and RSP, can lead to deficits in sensory-guided actions, such as reaching, eye movements for object tracking, and spatial navigation^45–48^. These findings further suggest the existence of neural circuits dedicated to transmitting sensorimotor error signals to the motor system. Although previous research has in fact identified circuits for computing prediction errors in the early sensory cortex^7,8,49^, understanding how error signals from V1 update predictions in frontal motor areas has been challenging due to the sparse axonal outputs from V1 to the frontal cortex (Fig. S17). By analyzing large-scale connectome data^50,51^, we found that axonal projections to the ACC from posterior higher-order areas (PM, AM, and RSP) were denser than projection from V1 and from both lateromedial (LM) and anterolateral (AL) visual areas to the ACC (Fig. S17). Our results from the electrophysiological and anatomical analyses suggest that these cortical circuits underlie the hierarchical transmission of prediction error signals from the PM, AM, and RSP to the ACC. Furthermore, axonal projections from the ACC to the posterior medial areas (PM, AM, and RSP) were also denser than those from the ACC to V1 and lateral higher-order visual areas (LM and AL) (Fig. S18). Notably, we also observed that visuomotor mismatch responses in PM and AM differed substantially from those in RL and AL, suggesting that anatomical pathway(s) from the ACC to PM, AM, and RSP convey top-down prediction signals, consistent with functional and anatomical investigations indicating the presence of two substreams in the dorsal visual pathway: a dorso–dorsal stream and a ventro–dorsal stream^17^. Collectively, these observed patterns of bidirectional anatomical connections provide additional support for the validity of hierarchical predictive coding in the cortex^2,5,19,52^.

### Dynamic Bayesian inference by prediction error signal

In sensory processing, it is essential to identify the source of sensory inputs. However, sensory signals processed in cortical circuits inherently contain noise introduced by transduction in the sensory organs and subsequent processing by neural circuits. In addition, the same physical information at the level of the sensory organs can convey different meanings depending on the context of the self and the environment. Consequently, the ability of the brain to “filter” and interpret sensory signals using dynamic, internally generated models is essential for producing a stable representation of the external environment^3^. In this study, we describe a SSM that generates error signals closely resembling those measured in mouse cortical neurons. Single-unit recordings revealed that the mismatch response patterns of individual neurons are more diverse than previously reported using Ca^2+^ imaging, in part due to greater temporal resolution. To characterize neuronal responses to mismatches, we focused on a parameter denoted *Q* representing state noise that provides a measure of adaptability to prediction errors in individual neurons. Previous models have included arithmetic subtraction scheme in which the observed visual flow is subtracted from the efference copy of locomotion-related signals to encode visuomotor prediction errors^19,34^. However, our analysis, which adapted the SSM to spiking activity, suggests that the neural response patterns to sensorimotor mismatch events may be better explained by a sequential probabilistic inference framework, such as the dynamic Bayesian inference framework^1,53,54^, where perception (defined as a posterior probability) is formed by updating predictions (defined as a prior probability) with observations. These findings highlight the possibility that flexible computation based on Bayesian inference allows animals to respond rapidly and precisely to dynamic environments by processing prediction errors.

### Evaluation of prediction error signals by response to mismatch events

Neural responses to visuomotor mismatch can be influenced by numerous factors, including visual features, concomitant motor activity, and prediction errors. Previous studies have shown that mismatch responses cannot be explained by motor or visual components alone, by using visual stimuli with matched visual input components (open-loop playback halt) to assess response properties, or by restricting postnatal visual experience to prevent the formation of visuomotor predictions in mice^49^. The present study was also designed to exclude potential confounding factors associated with visuomotor mismatch stimuli. Like previous studies, open-loop playback of visual flow cessation did not evoke large mismatch responses in the cortex (Fig. 1)^31,49^. However, while the playback paradigm can reproduce the visual inputs in closed-loop mismatch events, possible effects caused by differences in locomotion-dependent signals remain. The semi-closed-loop paradigm (SCL-VR) developed in the present study allowed us to assess the predictability dependence of mismatch responses under nearly identical visual inputs and locomotion states (Fig. 2). Notably, predictability established by prior experience to self-induced mismatch events affected both neural and pupil responses (Fig. 2). Pupil dilation, which regulates light input to the retina, occurred ∼0.25 s after the neural population mismatch response. Thus, it is plausible that the pupil dilation reflects the mouse’s perception or surprise to prediction errors rather than the cause of the mismatch responses. Other factors, such as locomotion, facial movements, eye movements, blinking, and the position and scene of the mouse in the VR, are ineffective explanatory variables for predicting the mismatch response. Rather, visuomotor prediction errors computed by our proposed SSM were a more plausible variable for explaining the neuronal responses (Fig. 4). These results support the notion that mismatch responses are caused by prediction errors^55^.

### Limitations of this study

The present study reveals a hierarchical processing stream of visuomotor prediction errors in the dorsal visual stream. Despite the use of wide-field imaging, however, these findings do not exclude the existence of other pathways transmitting sensory prediction errors, including subcortical streams. Neurons in dorso-dorsal visual areas, such as the PM, project axons not only to the ACC but also to the secondary motor cortex, RSP, lateral posterior nucleus, and striatum^18^. These regions are involved in visual information processing, motor execution, and learning, suggesting that prediction errors may also be transmitted to these regions. Moreover, previous studies have indicated that subcortical areas, such as the lateral posterior nucleus and superior colliculus, modulate sensory processing in the visual cortex during movement and thus likely receive error signals^56,57^. Depending on the nature of the internal or environmental context, error signals can be routed to brain regions that generate prediction signals. Elucidating the interaction between these cortico-subcortical and cortico-cortical circuit mechanisms will provide deeper insights into the neural basis of integrating sensory inputs with motor outputs for adaptive sensory perception and actions.

## Conclusion

This study provides experimental evidence for hierarchical cortico–cortical interactions underlying predictive coding. We demonstrated hierarchical propagation of bottom-up visuomotor prediction error signals from V1 to medial higher-order cortical areas, and to the frontal cortex, a potential source of predictions. This sensorimotor prediction error circuit can serve as a mechanism for updating the internal model for sensory perception and sensorimotor integration. Predictive coding is a computational framework that explains sensory perception and sensorimotor integration but is also applicable to understanding psychiatric and neurological disorders as an imbalance between predictions and actual inputs^58^. Understanding predictive processing at the cellular and circuit levels, in addition to the theoretical level, could contribute to the development of diagnostics and treatments for such disorders.

## Methods

### Animals

All animal procedures were conducted according to institutional and national guidelines, and were approved by the Animal Care and Use Committee of Nagoya University. All efforts were made to reduce the number of mice used and minimize their suffering and pain. C57BL/6J mice were purchased from Nihon SLC. Thy1-GCaMP6f (GP5.17) mice were purchased from the Jackson Laboratory (JAX025393)^37^. Mice were maintained in a temperature-controlled room (24 °C ± 1 °C) under a 12 h light/dark cycle with *ad libitum* access to food and water. Mice aged 11−28 weeks were used for the virtual reality (VR) experiments because of the body size needed for adaptation to head-fixed VR. All mice were reared under the normal light/dark cycle to evaluate the effect of their experiences in the VR.

### AAV production

All plasmids were constructed as previously described^59–61^, sequence-verified, and tested by transient expression in HEK293T cells before virus production. The AAVs were generated in HEK293T cells and purified by density gradient centrifugation, as previously described^60,61^. Genome titers of the AAVs were quantified by qPCR for the WPRE sequence. The titers of AAV-PHP.eB-hSyn-jGCaMP7f and AAV-PHP.eB-hSyn-GFP were 3.25 × 10^13^ and 2.96 × 10^13^ vg/mL, respectively. The AAV9-CAG-DIO-GtACR2-FusionRed-Kv2.1, AAV2retro-ESyn-iCre, and AAV2retro-hSyn-tdTomato were 1.10 × 10^13^, 1.42 × 10^13^, and 1.02 × 10^10^ vg/mL, respectively. Virus aliquots were stored at −80 °C until use.

### G-deleted rabies virus production

A G-deleted rabies viral vector encoding green fluorescent protein (RVΔG-GFP) was produced as previously described^59–62^. In brief, RVΔG-GFP was recovered by transfection of B7GG cells with the rabies viral genome vector pSAD-B19ΔG-GFP, pcDNA-SAD-B19N, pcDNA-SAD-B19P, pcDNA-SAD-B19L, and pcDNA-SAD-B19G. For *in vivo* injection, RVΔG-GFP was amplified in ten 15-cm dishes in 3% CO_2_ at 35 °C, filtered using a 0.45-μm filter, concentrated by two rounds of ultracentrifugation, and titrated with HEK-293T cells. The titer of RVΔG-GFP was 1.0 − 1.2 × 10^9^ infectious units/mL. Virus aliquots were stored at −80 °C until use.

### Retro-orbital virus injection and surgery

The AAV-PHP.eB-hSyn-jGCaMP7f was injected at 100 µL into the retro-orbital sinus of mice using a 30-gauge needle under deep anesthesia with a mixture of medetomidine hydrochloride (0.75 mg/kg; Nihon Zenyaku), midazolam (4 mg/kg; Sandoz), and butorphanol tartrate (5 mg/kg; Meiji Seika). Skull skin and membrane tissue on the skull were carefully removed, and the skull surface was covered with clear dental cement (204610402CL, Sun Medical) 14−20 days after virus injection. A custom-made metal head-plate was implanted onto the skull for stable head fixation. The visibility of GCaMP- or GFP-derived green fluorescence was verified under a fluorescent macroscope (M165FC, Leica). During surgery, the eyes were covered with ofloxacin ointment (0.3%) to prevent dry eye and unexpected injuries, and body temperature was maintained with a heating pad. After surgery, mice were injected with atipamezole hydrochloride solution (0.75 mg/kg; Meiji Seika) for rapid recovery from the effect of medetomidine hydrochloride.

### Virus injection for retrograde or anterograde labelling

Viral solution was injected stereotaxically into the mouse brain, as described previously^62^. Briefly, 8-week-old mice were anesthetized using the same procedure as that for retro-orbital injection. Mice were head-fixed on a stereotaxic apparatus (David Kopf Instruments). RVΔG-GFP (400 nL for each location) was injected into RSPa (AP: −1.5 mm, ML: 0.2–0.3 mm, depth: 0.3 and 0.6 mm from bregma) or RSPp (AP: −3.0 mm, ML: −0.5 mm, depth: 0.5 and 1.0 mm from bregma) at 100 nL/min with pulsed air pressure. AAV2retro-hSyn-tdTomato (400 nL) was injected into the ACC region (AP: -0.4 mm, ML: 0.5 mm, depth: 1.0 mm) according to the same surgical procedure of RVΔG injection. Injection sites of AAV9-CAG-axonGCaMP6s (200−300 nL) were determined by with intrinsic signal optical imaging in two of the four mice per treatment group, while injection sites in the other two mice per group were defined by distance from their bregma.

### Immunostaining

Mice were anesthetized and transcardially perfused with 50 mL of phosphate buffered saline (PBS) followed by 50 mL of 4% paraformaldehyde (PFA, Nacalai) in PBS. Brains were post-fixed with 4% PFA in PBS, 15% sucrose in PBS overnight, and then with 30% sucrose in PBS. The brains were subjected to histological analysis as described previously^60,62^. Cryoprotected samples were sectioned at a thickness of 40 µm on a freezing microtome (REM-710, Yamato Kohki). The sections were rinsed with PBS, blocked for 1−2 h in blocking solution (50% Blocking One (Nacalai, 03953-95) in 0.1% Triton X-100/PBS (PBST)), and then incubated overnight at 4℃ with primary antibodies: rabbit anti-GFP (1:3000, Abcam, ab6556), rat anti-GFP (1:1000, Nacalai, 04404-84), chicken anti-GFP (1:2000, Abcam, ab13970), rabbit anti-DsRed (1:1000, Clontech, 632496), goat anti-parvalbumin(1:500, Swant, PVG-213), rat anti-somatostatin (1:500, Millipore, MAB354), and rabbit anti-vasoactive intestinal peptide (1:400, ImmunoStar, 20077) in 5% Blocking One/0.1% PBST. After three washes with 0.01% PBST, the sections were incubated with Alexa Fluor 488-, 594-, 647-labelled donkey anti-rabbit IgG, anti-rat IgG, anti-goat (all 1:1000, Jackson, 711-545-152, 711-585-152, 711-605-152, 712-585-153, 712-605-153, 705-605-147) and DAPI (1:800, 1.25 μg/mll; Wako).

Images of immunofluorescence staining were acquired using a laser scanning confocal microscope (Zeiss, LSM800) with a Plan Apo 10× (Zeiss, NA 0.45) or Plan Apo 20× (Zeiss, NA 0.75) objective. The specificity of the primary antibodies was confirmed by typical staining patterns. None of the observed labeling was owing to nonspecific binding of secondary antibodies or autofluorescence in the fixed tissue because sections treated with secondary antibodies alone had no detectable signals. The acquired data were processed using ImageJ/Fiji for visualization.

### VR environment

Mice were head-fixed but free to run forward or backward on a treadmill. Three frameless 3/4-inch monitors (LP097QX1, LG) covered with antireflective film (TB-A18MFLKB, ELECOM) were used for visual stimulation. A 120-mm diameter urethane sponge was used as a treadmill stage. For two-photon Ca^2+^ imaging, backlight flickering was synchronized to the X-axis scan of the microscope to reduce light contamination. The renderings displayed on these monitors were precisely synchronized to a “Mosaic” function provided by the graphics card (Quadro P4000 or RTX-A2000, NVIDIA). Speed of the subject’s locomotion was monitored at the rotary encoder (360 pulses/cycle) connected to a microcontroller (Arduino Uno or Nano every, Arduino CC). The time series data of running speed were smoothed offline using a Savitzky−Golay filter (order = 3, frame length = 21). Visual feedback was provided by custom-written software using Open-GL API in MATLAB (MATLAB 2020a, MathWorks) or Python. MATLAB-based software was used in the wide-field imaging experiments. Python-based software was used in the two-photon imaging, optogenetic, and electrophysiological recording experiments. The latency of visual feedback was typically <50 ms (corresponding to <3 frames). The timing trigger for mismatch events was sent by the data acquisition board on the master PC (USB-6343, National Instruments) and received by the data acquisition board on the stimulus PC (USB-6002, National Instruments). For two-photon imaging experiments, the backlight LED of the monitors were synchronized to the resonant scanner turnaround points to prevent light contamination from fluorescence and visual feedback monitors. The TTL pulse for synchronization was generated by FPGA-based board (Analog discovery 2, Digilent)

For wide-field imaging data (Figs. 1, 2 and 5), the virtual environment simulated an infinite corridor with blurred random dot patterns. The patterns on the left and right walls were symmetrical, with no distinct landmarks. For cellular or axonal imaging, extracellular recording, and optogenetic experiments, mice were located on a virtual linear track with landmarks (200 cm/lap, four landmarks, Fig. 3). Visual flow stimuli were presented within the same VR environment. The flow speed corresponded to 31 cm/s locomotion for the 1.5 sec/trial. The rotary encoder speed, LED power, exposure timing, and mismatch or forced visual flow timing were recorded by the data acquisition interface of the master PC using a custom-written MATLAB application at a 20-kHz sampling rate. Mouse eye position, pupil size, and face were monitored by a machine vision camera equipped with an array of 940 nm IR-LEDs (AE-LED56V2, Akizuki-denshi), and a triacetate long-pass filter (cutoff wavelength = 820 nm, IR-82, Fujifilm). Room lights were off during the training and imaging sessions.

In the closed-loop VR condition, mismatches were triggered by a random timing generator program. Mice were habituated under the head-fixed VR condition for 1−2 h sessions for 3−5 days (Fig. 1) or 5– 9 days (Fig. 3–7) until they showed regular locomotion in the VR setup. Each imaging session under closed-loop VR also began with a habituation period of 15−30 min. For smooth head fixation, light anesthetization with (1.0%−1.5% isoflurane) was used at the beginning of the first habituation session.

In the semi-closed-loop VR (SCL-VR) condition, mismatch events were triggered by the stimulus PC in real-time when mouse locomotion speed exceeded a defined threshold (>12.0 cm/s, refractory period = 4.0 s). The logic flow of visual feedback in the SCL-VR is represented in Figure 2A. Six mice were habituated for 5 consecutive days (Fig. 2), and three of these were subsequently trained under the SCL-VR condition to experience predictable visuomotor mismatches (ME group), while the remaining three were trained under the regular closed-loop VR condition as predictable mismatch-inexperienced mice (NE group). The NE group mice were also habituated under the SCL-VR for an additional four days before the second imaging session (Fig. S9). The duration of habituation was 0.5 h for the first day and 1.5 h for days 2−5. On the first day of the habituation session, mice were habituated to darkness and treadmill running with head fixation.

Drifting grating stimuli and open-loop playback stimuli were presented on the same monitor as the VR using psychopy. Eight-direction rectangular gratings were presented 10 times for each direction (duration = 1.5 s, intertrial interval = 2.0 s). For open loop playback (playback-halt stimuli), screen captured video (captured in closed-loop mismatch imaging block) were presented using the “Movie stim” method of psychopy.

### Wide-field Ca^2+^ imaging

Calcium-dependent fluorescent signals from jGCaMP7f were acquired using a customized macroscope (THT, Brainvision) with a tandem lens design (a pair of Plan Apo 1×, WD = 61.5 mm, Leica) and epi-illumination system (see Fig. S2). An alternating excitation method was used to filter the calcium-independent fluctuation from the Ca^2+^ signal. Alternating blue (M470L4, Thorlabs) and violet (M405L3, Thorlabs) LEDs provided the excitation, which was filtered by additional band-pass filters (blue: #86-352, Edmund; violet: FBH400-40, Thorlabs). The excitation light was spatially equalized in the imaging area by a glass diffusion filter (DGUV10-600, Thorlabs). The average excitation LED power delivered to the surface was <5.0 mW. Fluorescent emission was passed through a dichroic mirror (FF495-Di03, Semrock) and a combination of long- and short-pass filters (FEL0500 and FESH0650, Thorlabs) to an sCMOS camera (ORCA-Fusion, Hamamatsu Photonics). The timing of the excitation light was controlled by a global exposure timing signal from the camera. The timing signal to switch the LED was processed with an FPGA-based logic circuit (Analog Discovery 2) and binary-counter IC (TC4520BP, Toshiba) in real time. In addition, the timing and power of the light source were monitored by a high-sensitivity photodetector (PDA10A2, Thorlabs).

All fluorescent emission images were recorded using HCImage Live software (Hamamatsu Photonics). Time-lapse images were acquired at 576 × 576 pixels (4 × 4 pixels binning). Single-shot high-resolution images (2304 × 2304 pixels) were also acquired to identify anatomical landmarks at the end of the imaging session. The sampling rate of each wavelength image was 20.0 Hz, and the practical exposure time (the duration of LED illumination per single frame) was 13.5 ms. The field of view was blocked from the light of the VR monitor by custom 3D-printed poly-lactic acid parts painted with matt black lacquer composition to prevent the contamination of the fluorescent signal and visual stimuli. All imaging sessions were shorter than 2.5 h. The surface of the coated skull was covered with a silicone material after the imaging session to maintain the visibility of the fluorescence. Only mice that showed regular locomotion on the treadmill were tested in further experiments.

### Extracellular recording

All extracellular recordings were performed with 32-channel silicon probes (A1x32-Poly2-10mm-50s-177 or A1x32-Poly3-5mm-25s-177, NeuroNexus) and an Open-Ephys Acquisition Board or a 384-channel Neuropixels 1.0 probe connected to a PXI-based system. Open-Ephys GUI was used for acquisition software^63^. Recorded signals were further processed using a spike-interface package (filtered at 600−6000 Hz for NeuroNexus data or 300−10000 Hz for Neuropixels data), and a common median reference was used for removing artifacts. The Neuropixels probe was inserted 3.2–3.85 mm beneath the cortical surface to measure responses in the LGN and V1 (Fig. S10); the hippocampus traversed by the probe were likewise recorded. Spike sorting was performed using Kilosort (version 2.0 for 32-channel NeuroNexus probe, version 2.5 for the Neuropixels probe)^64^. Intrinsic signal optical imaging was performed for all subjects to identify insertion points for V1 and PM. Insertion trajectory planning for the Neuropixels probe was based on Allen CCFv3. Accurate targeting was confirmed by post-hoc validation with DiI staining all mice. Detected clusters were manually inspected and curated using the Phy package. Mice were trained for at least one session in the dark in a head-fixed position (Track 3, Fig. 3), and five sessions in the closed-loop VR (Track1, Fig. 3). Two to four recordings were performed per mouse. Mice were settled in at least two re-habituation sessions (at Track 1) after each recording session. Five mice (recorded by the NeuroNexus probe) were recorded in Track 1−3. Four mice (recorded by the Neuropixels 1.0 probe) were recorded in Track 1 and 3.

### Two-photon Ca^2+^ imaging

Two-photon Ca^2+^ imaging experiments were performed as previously described^65^ with modifications using a dual-plane imaging microscope. The water-immersion objective lens (CFI75 LWD 16× W, Nikon, NA0.8) and femtosecond laser (Insight DeepSee+, Spectra-Physics) were equipped for two-photon imaging. To monitor the neural activity without animal tilting, the objective was mounted on a custom-built extension and rotation adapter. A spatial light modulator and electrically tunable lens were used to image two planes at 30.1 Hz total frame rate (∼15.05 Hz per plane) using 920 nm excitation. A head plate was implanted on Thy-1 GCaMP6f mice in the same manner as for the wide-field imaging. AAV2retro-hSyn-tdTomato was injected into the ACC to label ACC-projecting neurons. A layered glass imaging window was implanted after virus injection surgery. One to three recording sessions were performed per mouse (at Track-1 in Fig. 3). At least two re-habituation sessions in the complete closed-loop VR environment were performed between each recording session.

Axon bouton imaging was performed by single plane with faster frame rate (∼30.14 Hz). Non-rigid image registration and preprocessing were performed using suite2p^66^. Detected ROIs were manually selected to reject false positive ROIs. ROIs that have highly correlated activity during an imaging session (Pearson’s *r* > 0.85) were merged to prevent overcounting of boutons from the same neurons.

### Pathway-specific inhibition

Pathway-specific inhibition during mismatch periods was performed by optogenetics using GtACR2 with 488-nm stimulation (Coherent, OBIS 488). The total stimulation laser power was set at 40 mW (∼1.45 mW/mm^2^). Following training in the VR, AAV2retro-ESyn-iCre (300 nL) was injected into the bilateral ACC (0.3–0.5 mm lateral and 0.4-0.5 mm posterior from bregma), and AAV9-CAG-DIO-GtACR2-FusionRed (500 nL) was injected into the anteromedial part of the bilateral PM (1.8 mm lateral and 3.0 mm posterior from bregma). Behavioral tests were conducted 3–4 weeks post-injection, followed by two recording sessions. In the first session, visuomotor mismatches were presented at random timing (10% probability per second) with laser stimulation on 50% of mismatch trials). In the second session, no visuomotor mismatch events were presented, but laser stimulation was delivered with random timing.

### Processing of wide-field imaging data

Acquired images were compressed to 288 × 288 pixels, and then motion correction was performed with the 405-nm excited images using efficient subpixel image registration algorithms. A single template image for registration was used for the motion correction procedure in each imaging session. Each pixel signal was processed by the linear regression method. The 405-nm channel was resampled at 40.0 Hz using the interpolation method, and then interpolated values were used for signal correction of the 470-nm channel. The processed signal was smoothed using a moving average filter (width: 50 ms). Thereafter, all frames were registered to the top-view image of ACCF (330 × 285 pixels, corresponding to 13.2 × 11.4 mm in the ACCF) using two anatomical landmarks along the midline (the center of the olfactory bulb and base of the RSP) according to a previous study with modifications (Fig. S3)^65^. The scale bar is absent in the registered images because of image transformation from the raw recorded image. Registration to the ACCF was confirmed for every mouse using both resting-state connectivity and retinotopic maps (Fig.1 and S3). A two-dimensional Gaussian filter was applied for pixel-based presentation of results (sigma = 2 pixels).

To quantify the change in dF/F of the mismatch, visual feedback, and locomotion onset events, we defined the baseline frame for each event trial. The baseline frame of each trial was the average of the frames during 0.50 s for random mismatch (Fig. 1 and 2), and 0.25 s for semi-closed-loop VR before event onset. Since visual flow speed and wall texture were symmetric, the images or dF/Fs of the left and right hemispheres were computed independently, then data from the same trials were averaged (the image of the left hemisphere was mirrored) in all experiments except for the pixel-based correlation analysis. For the ROI-based analysis, the size of the ROI was 4 × 4 pixels. Maps of *p*-values were calculated from one-sided Mann–Whitney U tests.

A bootstrap test was used to compare results from different groups or sessions. For the NE vs. ME group comparison (Fig. 2 and S8), the trial data from each group were resampled with a bootstrap procedure and two pseudo-distributions produced from trial-averaged responses. The *p*-value was then calculated by computing the X/N overlap ratio, where X indicates the overlapping samples between the two distributions, and N indicates the number of iterations (1000 times resampling performed). For the comparison across sessions with same mouse, a pseudo-distribution was produced by subtracting the data recorded from Rec.1 and Rec. 2, and then the ratio of trials in the distribution was used to calculate *p*-values. Results of the one-sided test for each timepoint and ROI are presented in Figure S9.

### Visuomotor state-space model

To investigate neural encoding of prediction error, we conducted the following two-step analysis: (1) model-based estimation of the visuomotor prediction error based on the observed behavioral data, and (2) encoding model-based identification of the relationship between the estimated prediction error signal and neural activities.

In the first step, we formulated the following state-space model (SSM) to reconstruct the unobservable prediction error from observed behavioral data.

State equation:

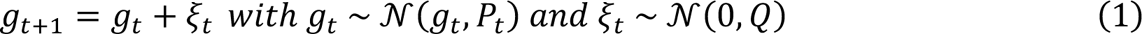

observation equation:

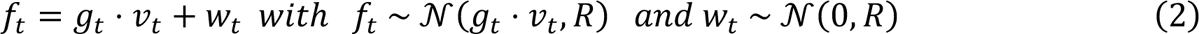

where, *f_t_*, *v_t_*, and *g_t_* stand for the observed visual flow speed, observed locomotion speed, and latent gain factor at time *t*, respectively. In this model, the state (i.e., latent gain factor) *g_t_* and the observation *f_t_* follow the Gaussian distribution 𝒩(*g_t_*, *p_t_*) and 𝒩(*g_t_* ⋅ *v_t_*, *R*). Note that *p_t_* is the covariance matrix of the state value’s distribution. The 𝜉_t_ and *w_t_* indicate the state and observation noise parameters, that follow the Gaussian distributions 𝒩(0, *Q*) and 𝒩(0, *R*), respectively. The *Q* and *R* are the fixed covariance parameters for the state and observation noise, respectively. As described in the experimental settings, we used mice that had learned that their locomotion speed on the treadmill is coupled to the speed of visual feedback flow in the VR environment. Therefore, we can assume that mice perform the locomotion task in the VR environment to satisfy that the visual flow *f_t_* matches the intended locomotion speed *v_t_* by adjusting the latent gain parameter *g_t_* generated as a motor command signal. Based on this assumption, we can predict the visual feedback speed *f_t_* based on the observed behavioral data (i.e., locomotion speed *v_t_*) by applying the above SSM. Thus, the extent of visuomotor mismatches caused by sudden changes in the actual visual feedback *f_obs,t_* can be quantified by calculating the prediction error *e_t_* between the exact and model-predicted speed of the visual flow feedback (i.e., *e_t_* = *f_obs,t_* − *f_t_*). To calculate such a prediction error, the unknown value of latent gain parameter *g_t_*should be estimated. Therefore, the following recursive Bayesian theorem was used to estimate the parameter *g_t_* based on given observational data *f_t_* and *v_t_*.

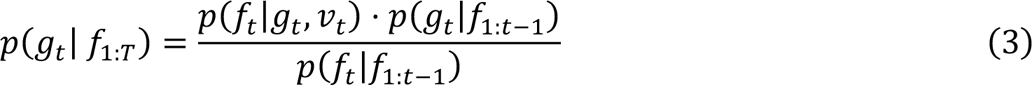

where, *f*_1:*T*_ stands for the observation set *f*_1:*T*_ = {*f*_1_, *f*_2_, …, *f_T_*}.. The *v_t_*indicates the observed value of the locomotion speed at time t, which is given as a known parameter in our proposed SSM. As mentioned above, because variables *g_t_* and *f_t_* both explain as a linear model following the Gaussian distributions *g_t_* ∼ 𝒩(*g_t_*, *p_t_*) and *f_t_* ∼ 𝒩(*g_t_* ⋅ *v_t_*, *R*), the above recursive Bayesian theorem can be solved by a linear Gaussian Filtering scheme (i.e., Kalman Filter; KF). By applying the KF, the recursive estimation rule of the parameter *g_t_* ∼ 𝒩(*g_t_*, *p_t_*), where the *g_t_*and *p_t_* stand for the mean and covariance of the Gaussian distribution, is given as the following two calculation steps:

[Prediction step]:

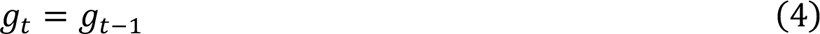

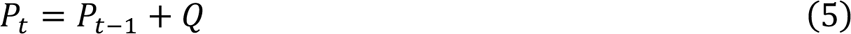

[Update step]:

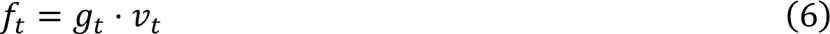

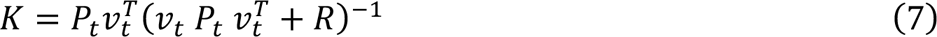

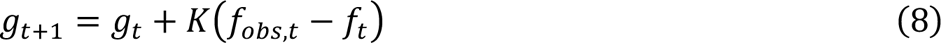

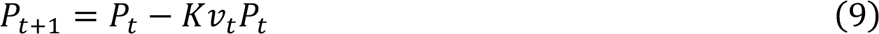

where, *f_obs,t_* and *f_t_* indicate the exact observation and model predicted value of visual flow speed at time *t*, respectively. The *v_t_* is locomotion speed that is given observed value. The *g_t_* is the latent gain factor at time *t*, which follows the Gaussian distributions 𝒩(*g_t_*, *p_t_*). *p_t_* is covariance of the variable *g_t_*. *K* stands for the Kalman gain.

Using the SSM and the KF-based estimation scheme, temporal changes in the visuomotor prediction error signal can be estimated from observations (in both *f_obs,t_* and *v_t_*). Here, the visuomotor prediction error is evaluated as: *e_t_* = *f_obs,t_* − *f_t_*. According to equation (8), the prediction accuracy of visual flow *f_t_* and resulting prediction error *e_t_*would be affected by the estimation of latent gain *g_t_*, which is parameterized by the extent of the state covariance *Q* . Therefore, when computing the visuomotor prediction error *e_t_*, the various predictions of *f_obs,t_* were evaluated under consideration with some different parameter settings for *Q*. In the following neural encoding model analysis, the neural response obtained from *in vivo* mouse brains was compared to the estimated prediction error *e_t_* with each condition of *Q*.

### Generalized linear model analysis

A generalized linear model (GLM) was used to estimate the time-dependent effects of experimentally measured variables. This GLM analysis was performed on data from mice that experienced all VR condition (Tracks 1–3). As explanatory variables, we used the mouse motion values (running speed, acceleration, facial movements, blinks, eye movements, and pupil diameter), VR-related values (position, visual flow speed, mismatch event timing, and track), and the SSM value described below. Data were further split by VR-laps, with odd laps forming the training set and even laps the test set. We used the “TweedieRegressor” class from the scikit-learn library for model fitting. The target distribution was set to “Poisson”, and the GLM link function was configured as “log”. To avoid overfitting, a weight vector was estimated by solving the penalized residual sum of squares using L2 regularization. Regression model hyperparameters were estimated through a grid-search of the training dataset.

Using the GLM, we evaluated the relationship between neural activities and the prediction error (*e_t_*) estimated by the above-mentioned SSM modelling scheme. For quantification of the individual neuronal sensitivity to prediction error information, two models for individual neurons were constructed: one fitted by a predictor matrix with full kernels (full model) and the other fitted by a matrix without a prediction error kernel (w/o SSM model). We then fit the models with each design matrix to predict the firing rates of individual neurons and calculated the explained variance (R^2^_Full_, R^2^ _w/o SSM_) of the full and partial models (Fig. 4). Finally, the neural encoding weight for prediction error was calculated as the difference in explained variance (ΔR^2^= R^2^_Full_-R^2^_w/o SSM_).

### Correlation map and GC analysis

To examine the information flow of the mismatch responses among cortical areas, pairwise correlations and Granger causality (GC) were computed between the activities of each ROI. GC analysis was conducted using the multivariate GC toolbox (pairwise conditional GC is termed “GC”). Neural signals before the mismatch onset (−1.5-0.0 s) were used to calculate the correlation map and GC magnitudes for closed-loop periods. The hyperparameter of GC estimation (autoregressive model order) was selected by AIC. Similarly, neural signals acquired during mismatch (0.0-1.5 s) were used to compute the correlation maps and the GC magnitudes for mismatch periods. The correlation coefficient was computed from the z-scored signal for each animal and the statistical significance of the correlation map was derived using the z-statistic value per pixel. Locomotion speed was used as a threshold to distinguish the time windows during which the subject was stationary or running in the dark (running, >6.0 cm/s; stationary, < 0.02 cm/s), while the speed range between these two thresholds was excluded.

### Face video analysis

A single CMOS camera (DMK33UX174 or DMK33UX273, The Imaging Source) affixed to the basement of VR system (MB3030D/M, Thorlabs) by a flexible arm to monitor facial and eye movements during experiments. The sampling rate of the video was approximately 20 Hz. All acquired frames were cropped to a resolution of 640 × 480 or 720 × 480 pixels to reduce post-processing. To evaluate the effects of face and eye movement, the positions of facial landmarks were extracted using DeepLabCut^67^. We defined the position of the eyelid (edges of the dorsal and ventral margins), pupil edge (dorsal, ventral, anterior, and posterior margins), and nose. The “resnet_50” configuration was used for the training of the pose estimation model. The error of extracting position was within ∼3 pixels. Blinking was detected by the rate of change in the distance between the upper and lower eyelids. Changes of > 25% of the distance during the mismatch period was defined as a blinking trial. Grooming and irregular behaviors were manually detected by the experimenter (R.F.T.). To extract the time series of motion energy in the face, pymoten^68^ was employed for the aforementioned movie data. Ten principal components calculated by PCA were used to reduce the dimensionality of the extracted signal.

### Dorsal cortex-wide imaging under the anesthetized condition

Functional mapping of cortical areas was performed after the recovery period (4−7 days) from head-plate implantation (see resting-state connectivity mapping and retinotopic mapping, below) and following the training and wide-field imaging sessions under VR. Briefly, mice were lightly anesthetized by isoflurane inhalation (1.0%–1.5%) during imaging. Body temperature was maintained with a heating pad. During the VR-imaging session, the face was recorded with a CMOS camera under infrared illumination to monitor mouse arousal. Sessions where the mouse showed distinct body movements were manually identified and excluded from the analysis. The isoflurane concentration was manually controlled based on body and facial movements as monitored by the facial camera. Visual stimuli were presented on the same display used for VR, and the treadmill setup was replaced by a stage with a heating pad and laboratory scissor jack.

### Resting-state connectivity mapping

Resting-state cortical Ca^2+^ dynamics were recorded from mice under light isoflurane inhalation (1.0%– 1.5%) without artificial sensory stimulation. Mice received retro-orbital injections of AAV-PHP.eB-hSyn-jGCaMP7f. The seed-based Pearson’s correlation map for each ROI was then calculated. R-squared values are used for visualization.

### Retinotopic mapping

Retinotopic mapping was performed to identify visual areas. Fluorescent signals were measured using macroscopic imaging through the skull or two-photon imaging through a glass-implanted window (Fig. 5 and 6). Drifting checkerboard stimuli flickering at 6 Hz were presented in four directions (0, 90, 180, and 270 degrees, T = 5 or 15 s). Typically, 30 cycles (for T = 5 s) or 19 cycles (for T = 15 s) of the stimuli were presented to mice. Acquired images were processed by phase-encoding methods to visualize the retinotopic map of the cortex.

Intrinsic signal optical imaging was also performed for retinotopic mapping. Injection sites for axon imaging or recording sites for extracellular electrophysiology were determined based on the identified visual areas. Imaging experiments were performed as previously described^62,69^. Mice were lightly anesthetized with isoflurane inhalation (0.8%−1.2%) in pure oxygen. Intrinsic signals from the cortex were acquired through the resin-coated skull using a CMOS camera under 625-nm LED illumination (M625L3, Thorlabs). The 5-Hz flickering and moving checkerboard horizontal or vertical stimuli were presented to mice. Phase-encoding analysis was performed for acquired images, and the visual field sign map was calculated.

### Connectivity analysis of the anatomical database

The Allen Mouse Brain Connectivity Atlas was used to quantify the anatomical connectivity among the dorsal cortex. The atlas consists of high-resolution images of axonal projections labelled with AAV injections into various locations. The normalized projection volume from 36 pooled experiments (RSPa-IDs: 100140949, 166458363, 516838033, 292172100, 272735744, 267661018, 177907082, 159097209, 526502961, 166271142, 182338356, 288264753, 100148142, 166269090, 159832064, 292124058, 267658040, 308721884, 184168193, 278179794, 591535205, 287601100, 166325321, 308027576; RSPp-IDs: 157711043, 298720191, 181860879, 538078619, 584895127, 298275548, 521264566, 592540591, 166054929, 272916915, 112424813, and 182467736) was used to quantify RSPa and RSPp output. For ease of interpretation, experiments using a combination of the Ai75 line and the CAV2-Cre virus vector were excluded from the analysis. Multiple experimental datasets were also aggregated for analysis of ACC input. The experiments considered relevant were identified based on their horizontal (AP and ML axis) distance from the defined ROI in macroscopic imaging data, specifically within a range of less than 500 µm from the center of the ROI (Fig. S17). For ACC output analysis, all experiments where the primary injection sites were identified as ACCd/ACCv were included (Fig. S18). These analyses were conducted using the Allen Software Development Kit within the Google Colaboratory environment.

## Acknowledgements

We thank members of the Osakada laboratory for their valuable discussions, Drs. K. Mizuseki, M. Ogawa, A.W. Ishikawa, R. Hira, M. Funamizu, Y. Osako, K. Amemori, and Mr. Y. Saito for discussions, Mr. Hanada, Nishimura, Kano, Kato, and Kobayashi (the Research Equipment Development Group, Technical Center of Nagoya University) for manufacturing instruments, and CARE members (Nagoya University) for the housing of animals. This work was supported by fundings by the Grants-in-Aid from JSPS (17H05562, 18H02706, and 21H05168 to F.O., 20K16464 to R.F.T.), PRESTO and CREST from JST (F.O.), and Brain/MINDS and CREST from AMED (F.O., JP19dm0207057, 19dm0207058).

## Author contributions

R.F.T. constructed all experimental setups, performed viral injections, collected and analyzed the imaging, extracellular recording, and optogenetics data, and wrote the manuscript. A.Y.S. collected the extracellular recording data and wrote the manuscript. K.N.I. performed viral injections and histological analysis and wrote the manuscript. H.Y. and N.H. contributed to model construction and manuscript writing. R.M. performed viral injections and collected and analyzed the behavioral and imaging data. R.U. performed viral injections and anatomical analysis. K.K. performed viral injections and animal surgery. R.K. and T.S. performed molecular cloning and virus production. K.I. constructed a dual-plane two-photon imaging system. F.O. wrote the manuscript and supervised the project.

## Competing interests

Authors declare that they have no competing interests.

### Data and materials availability

The datasets and viral plasmids generated in this study are available from the corresponding authors upon reasonable request. All data needed to evaluate the conclusions in this paper are included in the paper and/or the Supplementary Materials. Additional data related to this paper will be provided upon reasonable request.

## Extended data figures

**Figure S1.**
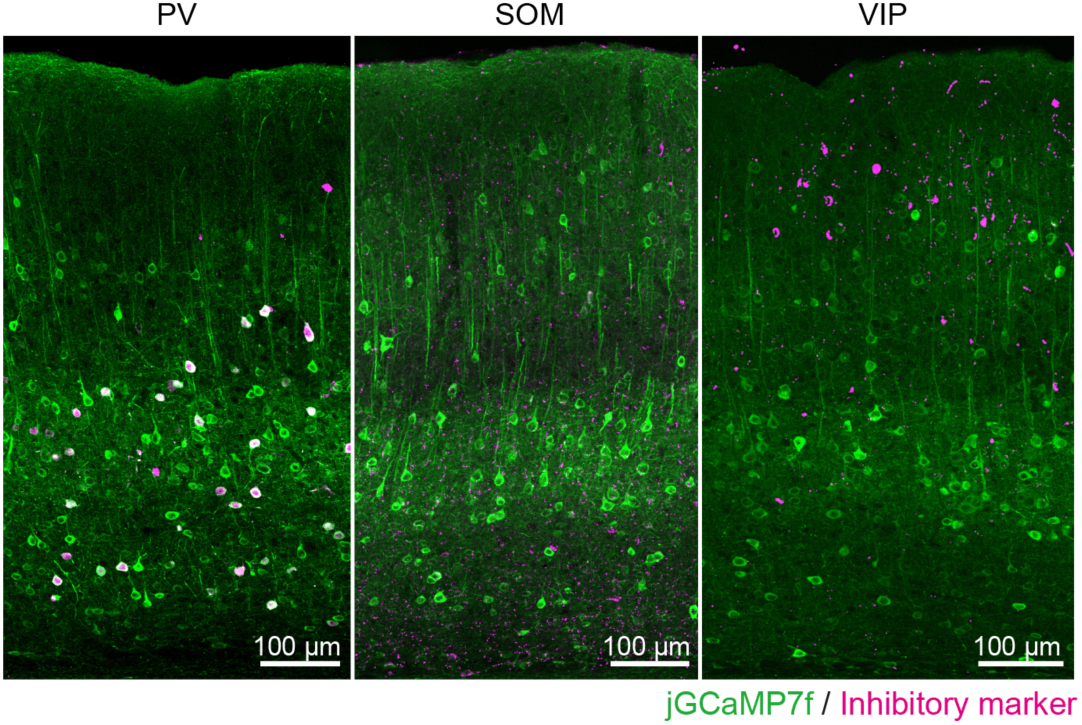
Excitatory neuron-biased expression of GCaMP, related to Figure 1. Confocal images of brain sections from AAV-PHP.eB-hSyn-jGCaMP7f-infected mice immunostained for the inhibitory neuron subtype markers parvalbumin (PV), somatostatin (SOM), and vasoactive intestinal peptide (VIP). Most jGCaMP7f-expressing neurons in the cortex showed typical morphological features of pyramidal cells and were distributed from superficial to deep layers. Only a few cells were positive for both jGCaMP7f and an inhibitory neuron marker.

**Figure S2.**
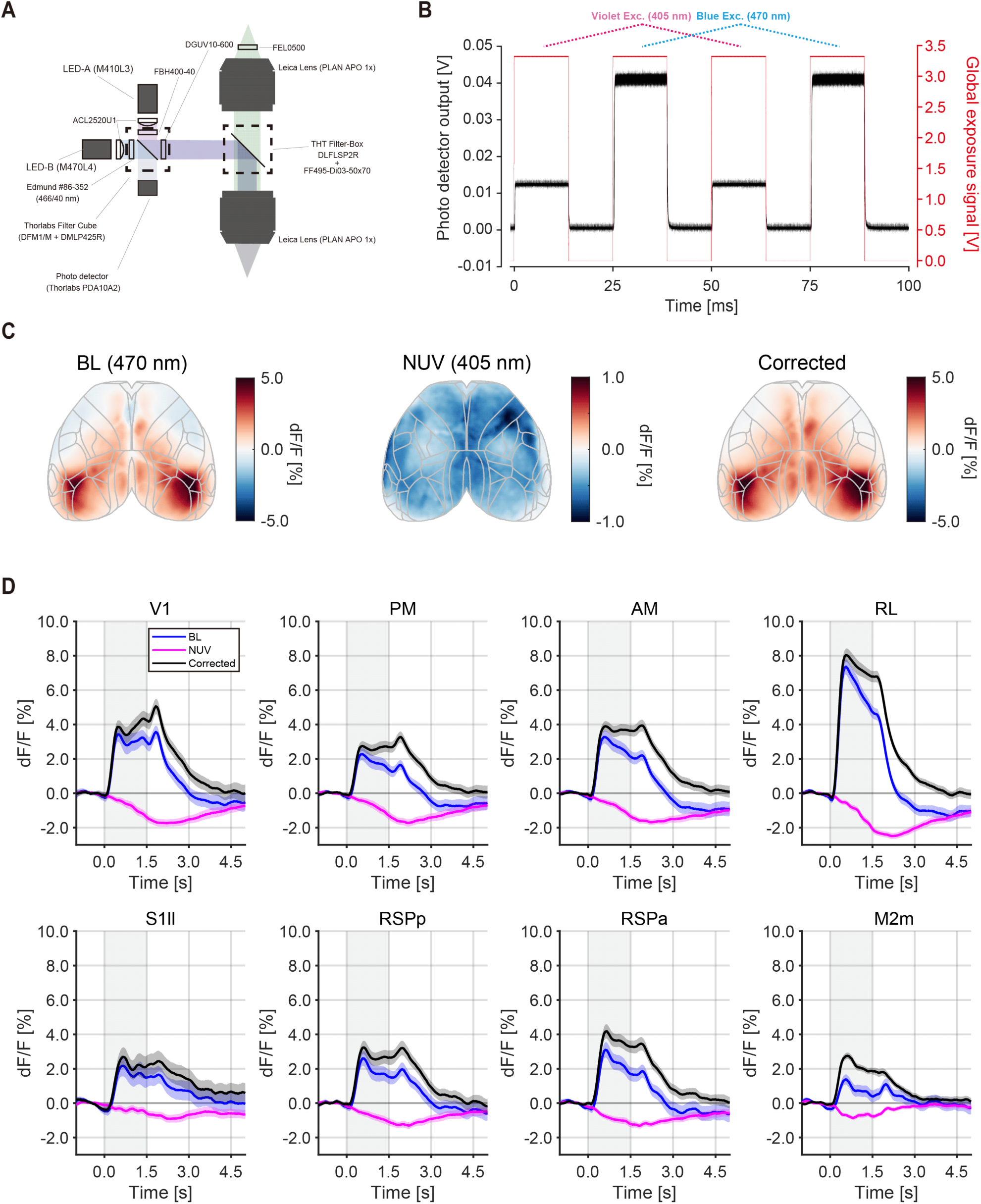
Signal-correction using near ultra-violet light, related to Figure 1. (A) Schematic of the optical fluorescent imaging system. (B) Latency of the global exposure strobe signal from the camera to the LED illumination at 405 nm and 470 nm. (C) Signal correction of neural response maps at 405 nm and 470 nm. Maps were generated by averaging the flow onset periods (0.0−1.0 s). (D) Neural responses of eight ROIs. Shaded regions represent the SEM. Gray shaded lines indicate visual flow periods.

**Figure S3.**
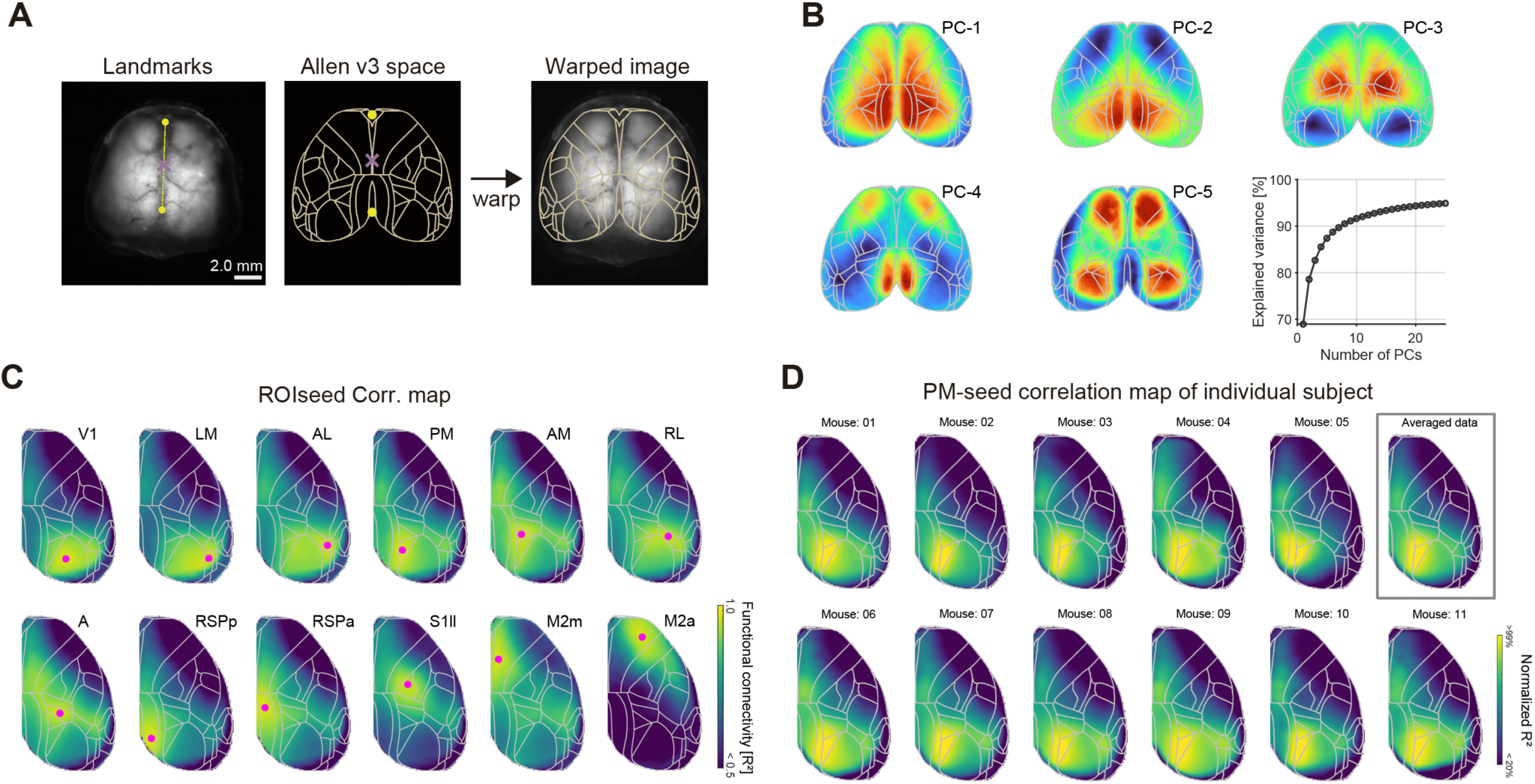
Registration of an individual brain image to the Allen common coordinate framework (ACCF), related to Figure 1. (A) Registration process from the acquired brain image to ACCF space. Left: Manually mapped landmarks on the preregistration fluorescence image of the living mouse brain (illuminated by 470 nm) and the areal boundary of the ACCF space. Right: Post-registration image overlaid with an area boundary. Gray outlines area parcellation from the ACCF. Yellow dots indicate two control points, the center of the olfactory bulb and the base of the retrosplenial cortex (RSP). Magenta crosses indicate the bregma. (B) Principal component analysis (PCA) of resting-state cortical activity. Weight maps of PCA (Component 1−5) derived from six mice. Right bottom: Cumulative explained variance by PCs. (C) Functional connectivity maps of the animals shown in Figure 1. Magenta dots indicate the center of the ROI for the seed-based correlation map. Maps are the average of the right and left hemispheres. V1, primary visual area; LM, lateromedial visual area; AL, anterolateral visual area; PM, posteromedial visual area; AM, anteromedial visual area; RL, rostrolateral visual area; S1ll, primary somatosensory area, lower limb; RSPp, posterior part of RSP; RSPa, anterior part of RSP; M2m, medial part of secondary motor area. (D) Seed-based (PM) correlation maps from mice used in Figure 1 and 2.

**Figure S4.**
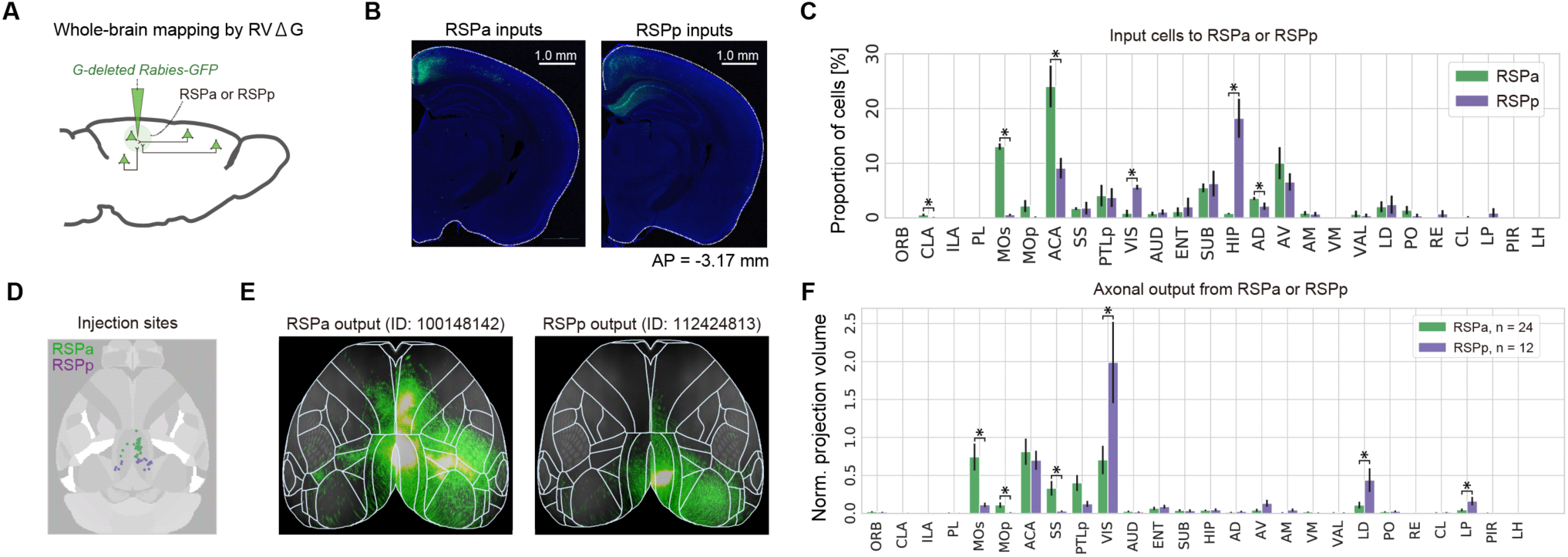
Anatomical differences in RSP inputs and outputs across the anterior-posterior axis, related to Figure 1. (A) Schematic of retrograde tracing from RSPa or RSPp. (B) Example images of cells retrogradely labelled with RVΔGFP. (C) Summary of inputs to RSPa or RSPp neurons. Asterisks show significant differences (*p* <0.05, by Mann–Whitney U-test). (D) Anterograde injection sites from the Allen mouse brain connectivity atlas. (E) Top view of axonal cortical output for example injection experiments (adapted from the Allen mouse brain connectivity atlas). Experiment IDs are indicated on the top. (F) Summary of inputs to RSPa or RSPp neurons. Normalized projection volume was used for the quantification (mean ± SEM). Asterisks show significant differences (*p* <0.05 by Mann–Whitney U-test).

**Figure S5.**
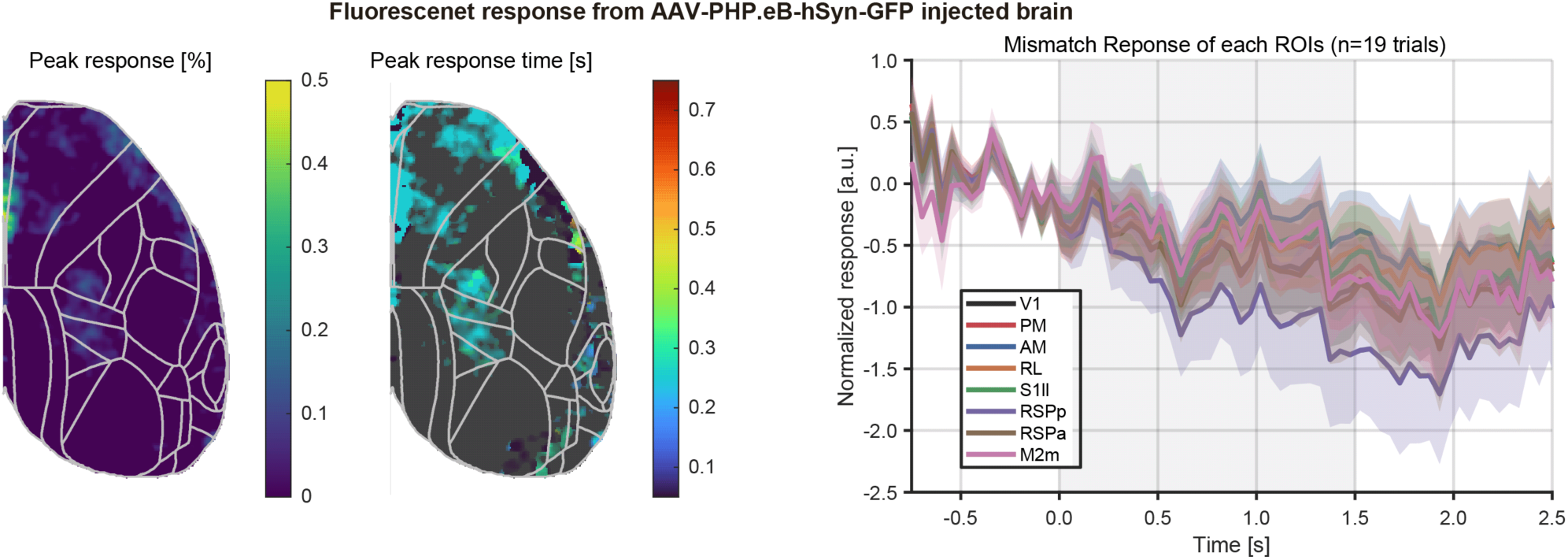
Ca^2+^ dependency of macroscopic mismatch responses, related to Figure 1. Left: Peak response amplitude map during mismatch periods (0.0−1.5 s from mismatch onset); Middle: Peak response time map during mismatch periods (0.0−1.5 s from mismatch onset); Right: Time course of mismatch responses in each ROI.

**Figure S6.**
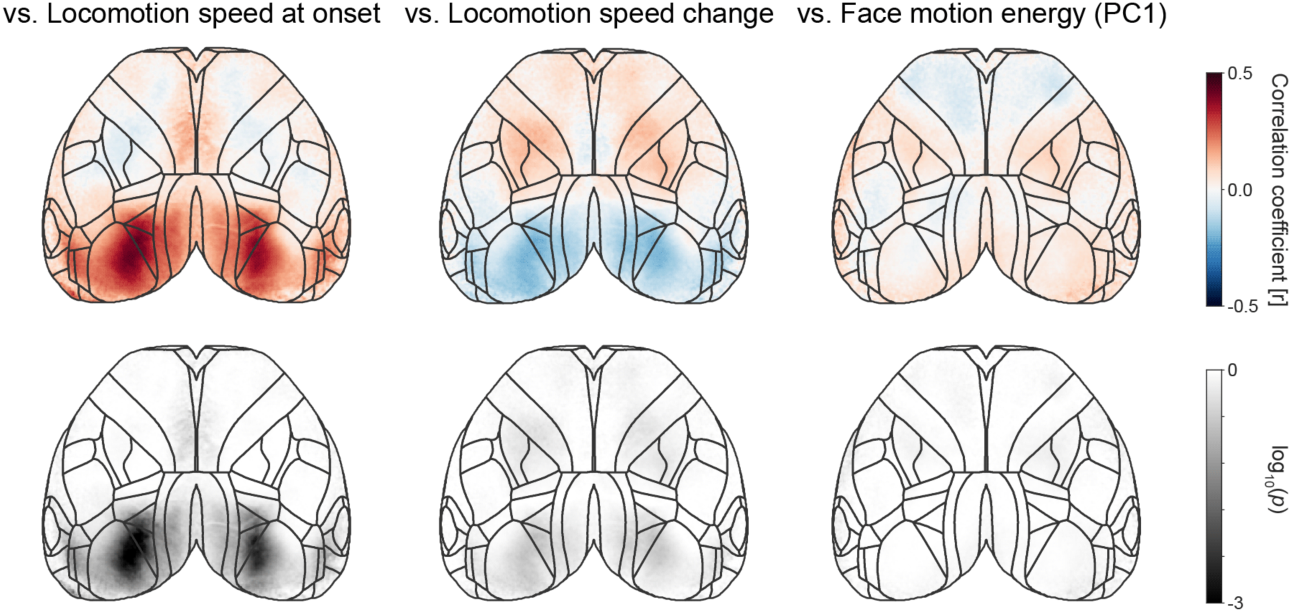
Explained trial variance of mismatch responses, related to Figure 1. Left: Pseudocolor map of Pearson’s linear correlation coefficient between locomotion speed at mismatch onset and neural response magnitude. Pearson’s correlation coefficient value (top) and *p*-value (bottom). Middle: Pseudocolor map of Pearson’s linear correlation coefficient between change in locomotion between pre- and post-mismatch onset and neural response magnitude. Pearson’s correlation coefficient value (top) and *p*-value (bottom). Right: Pseudocolor map of Pearson’s linear correlation coefficient between change in the first principal component (PC1) of the motion energy between pre- and post-mismatch onset and neural response magnitude. Pearson’s correlation coefficient value (top) and *p*-value (bottom). All results were computed from pooled data from nine mice.

**Figure S7.**
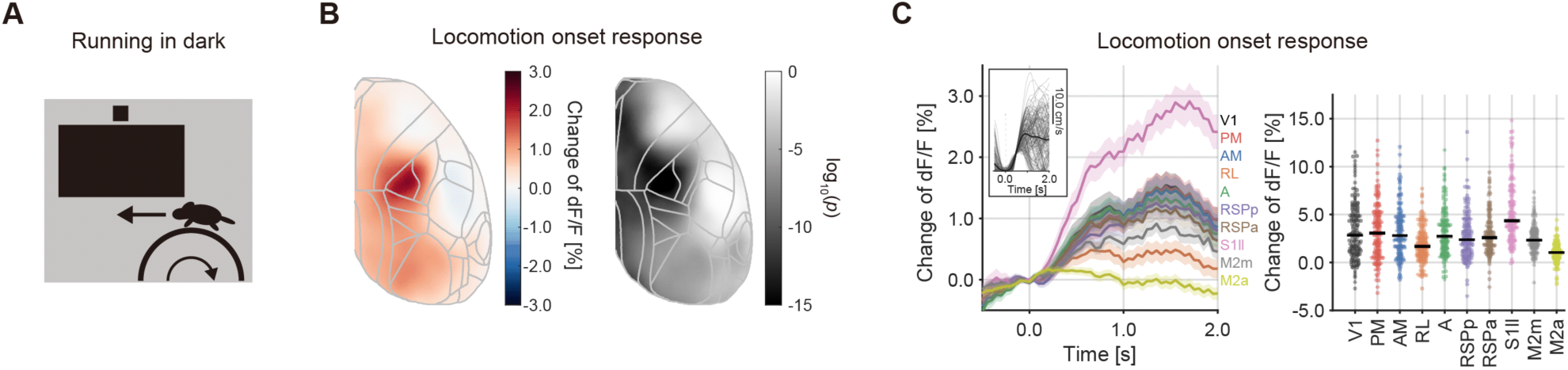
Macroscopic cortical activity evoked by locomotion, related to Figure 1. (A). Schematic of the experiments. (B). Locomotion onset response map. Left: Locomotion onset responses across the dorsal cortex (n = 5 mice). Right: Peak responses during the time window 0.0−1.0 s after the onset were quantified. (C). Quantification of locomotion onset responses in 10 cortical areas. Left: Locomotion onset responses in 10 cortical areas (n = 5 mice). Inset: Running speed of each trial. Right: Peak responses during the 0.0−1.0 s time window in each area.

**Figure S8.**
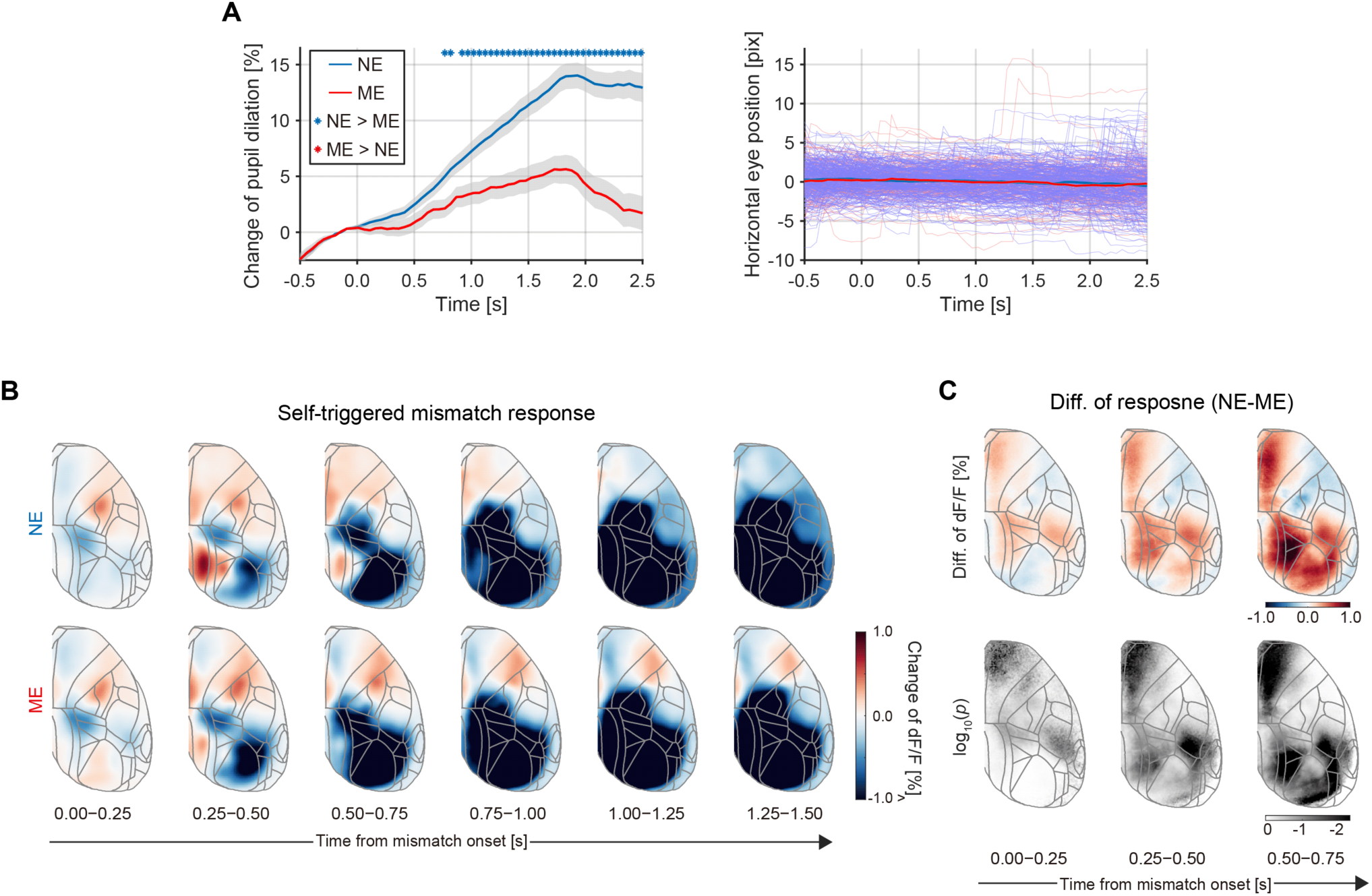
Time series of self-induced mismatch responses, related to Figure 2. (A). Time courses of pupil response and horizontal eye movements to mismatch events. (B). Trial-averaged neural responses to mismatch events (NE group: n = 268 trials from 3 mice; ME group: n = 156 trials from 3 mice). (C). Top: Difference in self-induced mismatch responses (NE minus ME). Bottom: Significant difference between NE and ME groups by one-sided Mann−Whitney U test.

**Figure S9.**
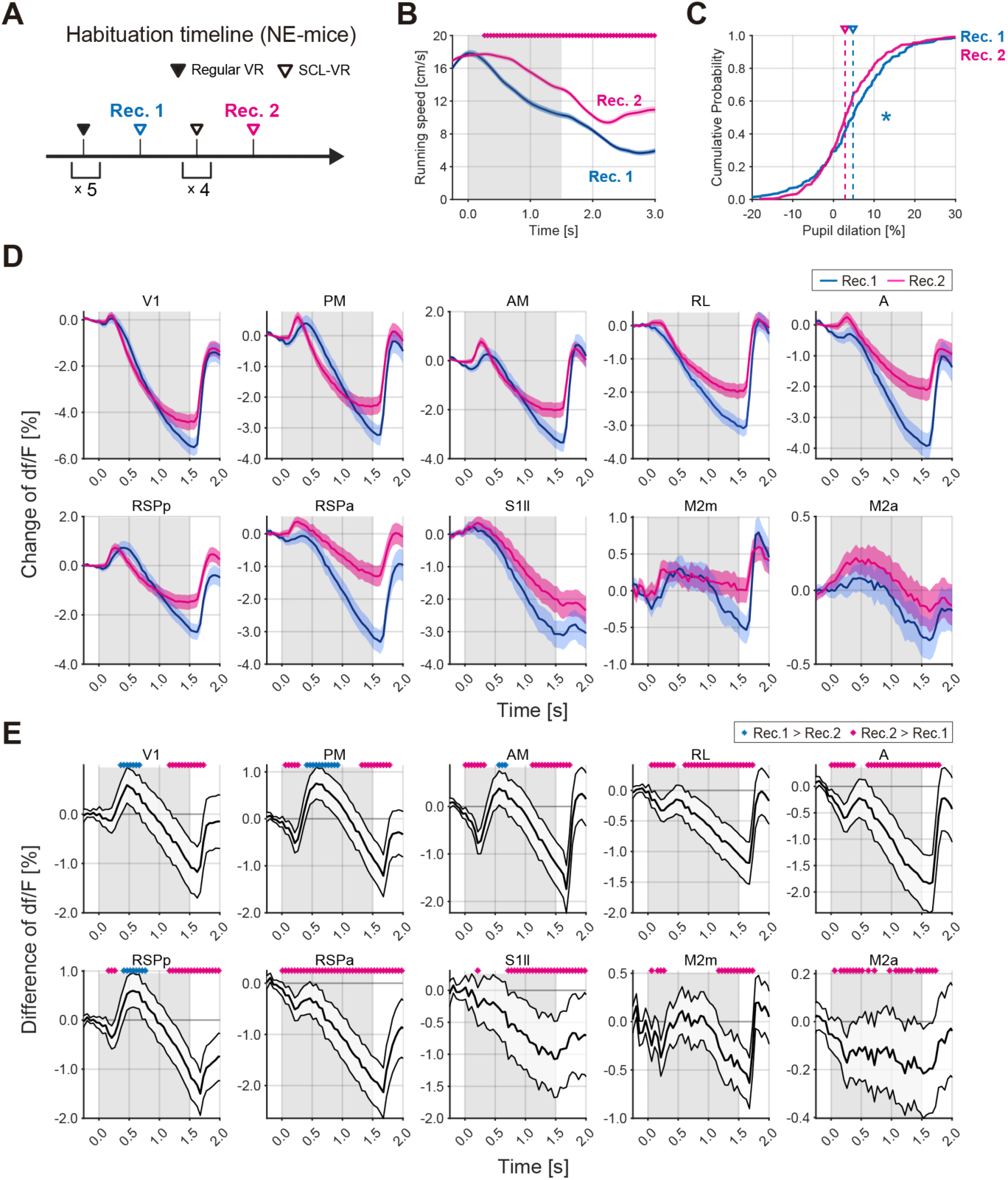
Reduction of mismatch responses by repeated exposure to self-induced mismatch, related to Figure 2. (A). Habituation history of NE mice (as in Fig. 2). (B). Locomotion speed of NE mice during self-induced mismatch events. The shaded region indicates the mismatch period (0.0−1.5 s). Line traces represent means ± SEM (blue: first recording session, magenta: second recording session). Dots show time bins with significant differences (*p* <0.01 by one-tailed unpaired *t*-test). (C). Cumulative distribution plot of pupil dilation responses during mismatch onset trials (0.0−1.0 s average). Dashed line indicates the median value for each group. The asterisk indicates a significant difference (*p* <0.05 by Mann–Whitney U-test). (D). Trial-averaged response traces from 10 cortical areas (ROIs). The shaded region indicates the mismatch period (0.0−1.5 s). Blue plots area results from the first recording session. Magenta plots are from the second recording session. Each plot is shown as a mean ± 95% confidence interval. (E). Difference in mismatch responses between recordings Rec. 1 and Rec. 2 for the same animal (subtraction was applied between the same animal’s data). Each plot is shown as a mean ± 95% confidence interval. Dots indicate time bins with statistical significance (*p* <0.05 by one-tailed bootstrap test). Dot color indicates the alternative hypothesis (blue: Rec. 1 > Rec. 2; magenta: Rec. 2 > Rec. 1).

**Figure S10.**
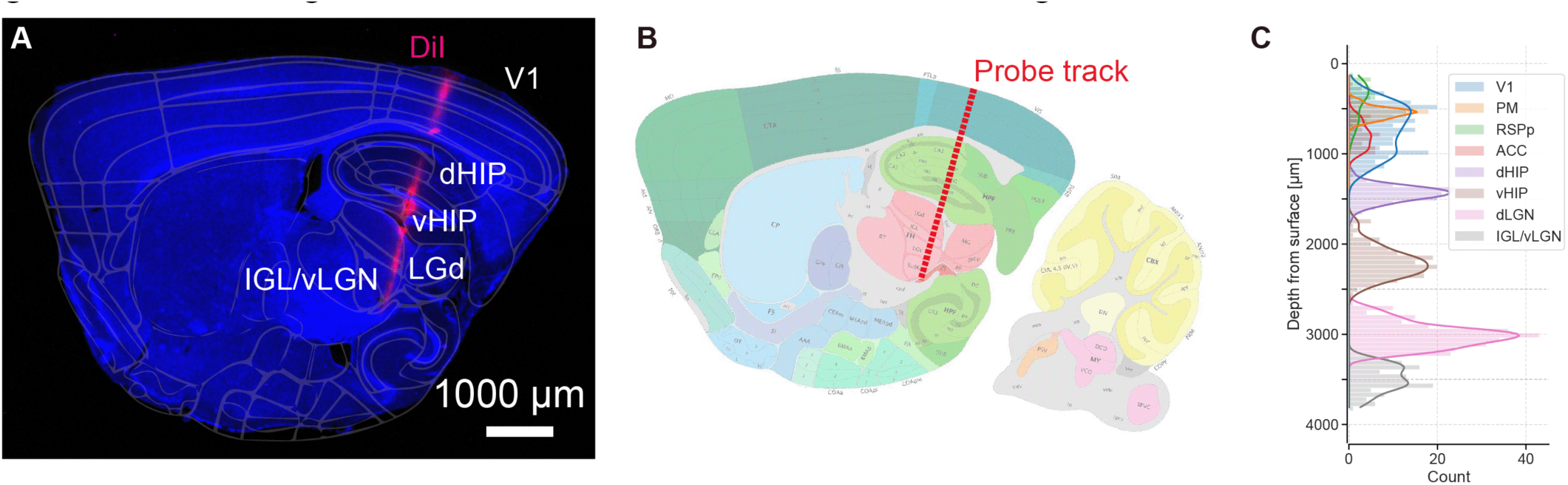
Recording site identification of extracellular recording, related to Figure 3. (A). Sagittal image of a mouse brain stained with DAPI (blue). The probe track was visualized with DiI (red). Areal boundaries estimated from the Allen brain atlas are overlaid. (B). Inferred probe track in the Allen CCFv3 atlas. (C). Distribution of isolated single units from the brain surface (pooled from 20 recordings).

**Figure S11.**
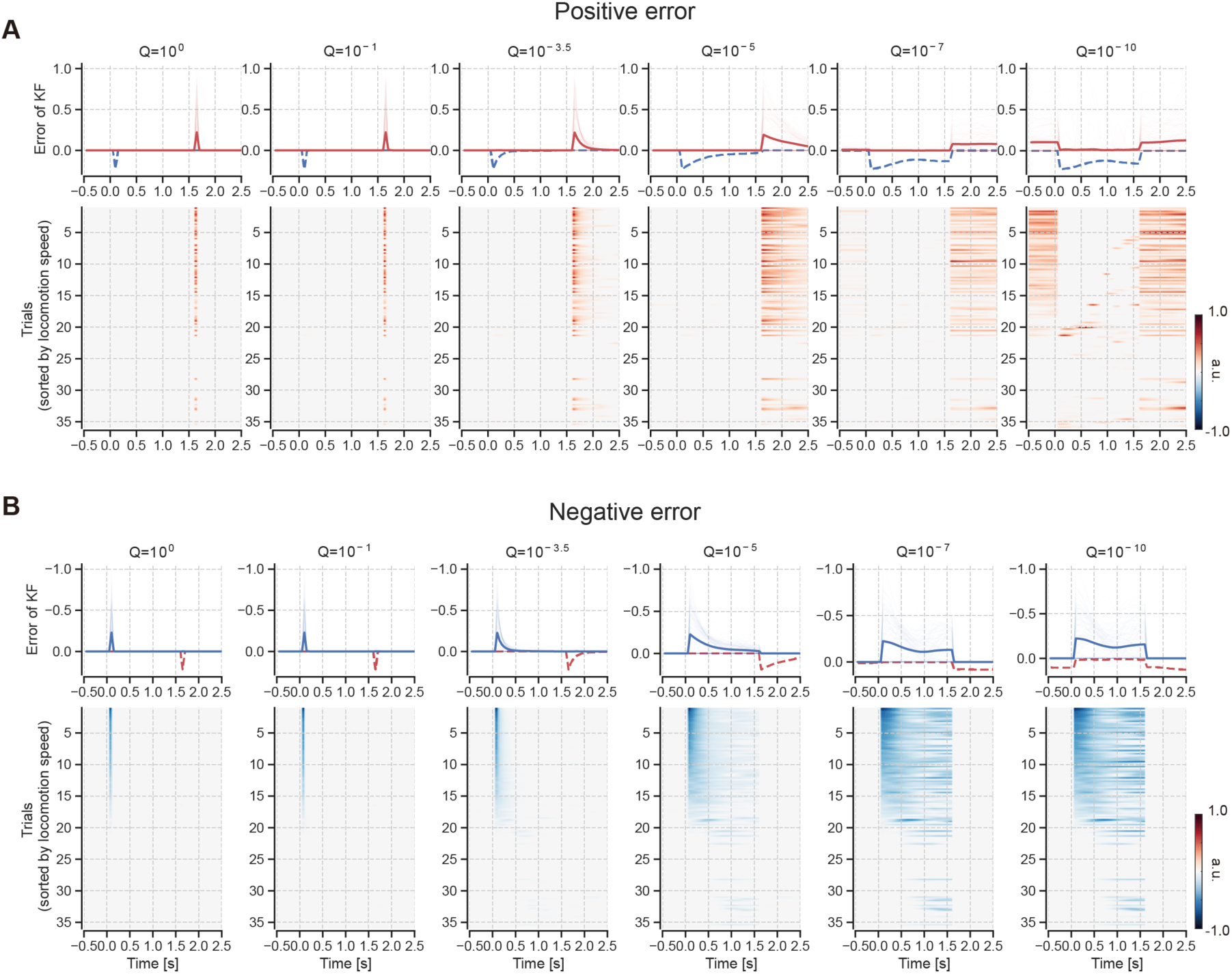
Visuomotor prediction error computed by state-space model (SSM) variation, related to Figure 4. (A) The SSM calculated the positive visuomotor prediction errors using actual experimental data obtained from an example session for input signals (locomotion speed and visual flow). Each column shows the distinct patterns of prediction errors generated by the SSM using distinct Q values. Bold line represents the trial-averaged calculated visuomotor prediction errors, while the thin red lines show the errors from each individual trial. The dashed blue line indicates negative error outputs from the corresponding Q models. (B) Same as in (a) but for negative error cases.

**Figure S12.**
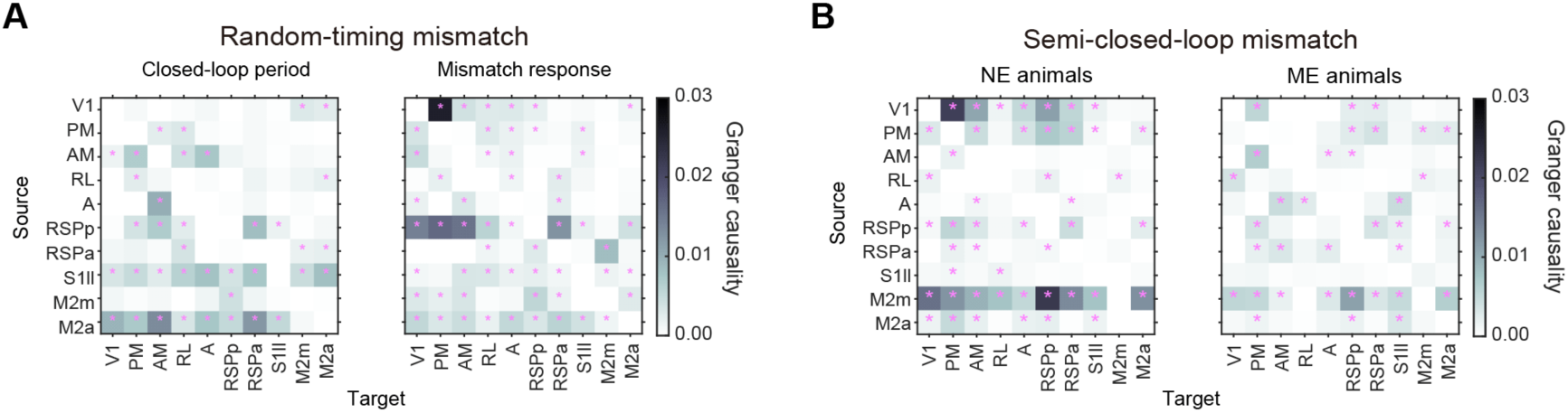
Granger causality results across dorsal cortical areas, related to Figure 5. (A). Granger causality between dorsal cortical areas during closed-loop and mismatch periods. Magenta dots represent statistical significance (*p* < 0.01 by permutation test). (B). Granger causality between dorsal cortical areas during semi-closed-loop mismatch periods for NE and ME groups.

**Figure S13.**
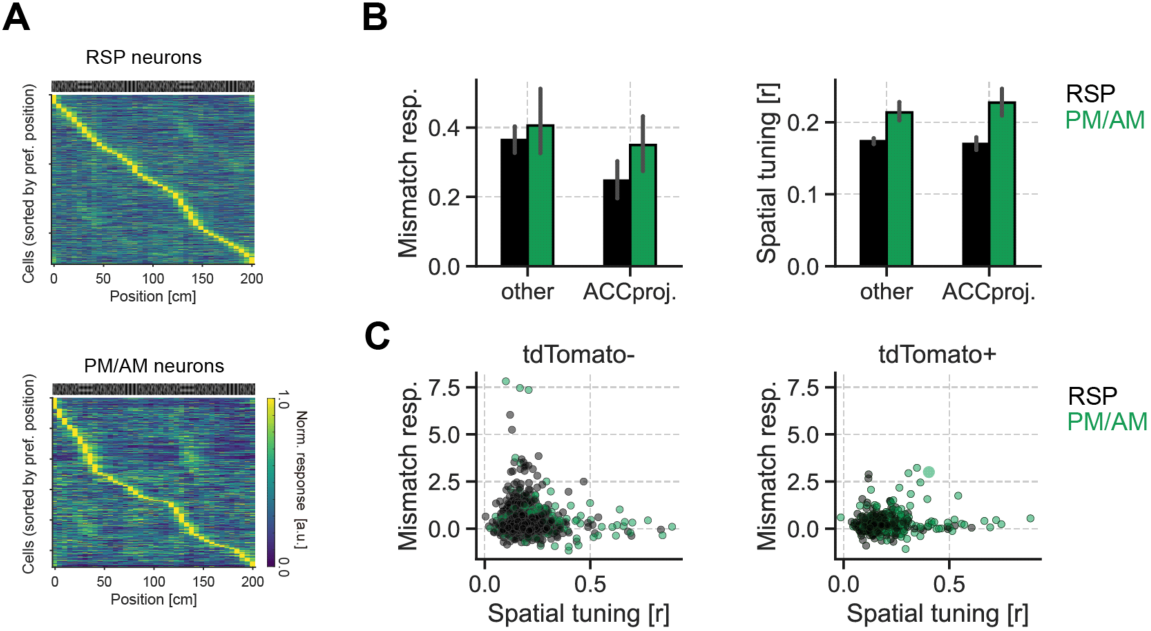
Tuning profiles of tdTomato^+/-^ neurons in RSP and PM/AM, related to Figure 6. (A) Sorted, trial-averaged position activity maps for all RSP and PM/AM cells. (B) Distributions of mismatch response magnitude (left) and spatial tuning (right). Error bars represent SEM. (C) Relationship between spatial tuning strength and mismatch response magnitude in tdTomato^-^ cells and tdTomato^+^ cells. Pearson’s correlation coefficients are all insignificant (*p* > 0.05).

**Figure S14.**
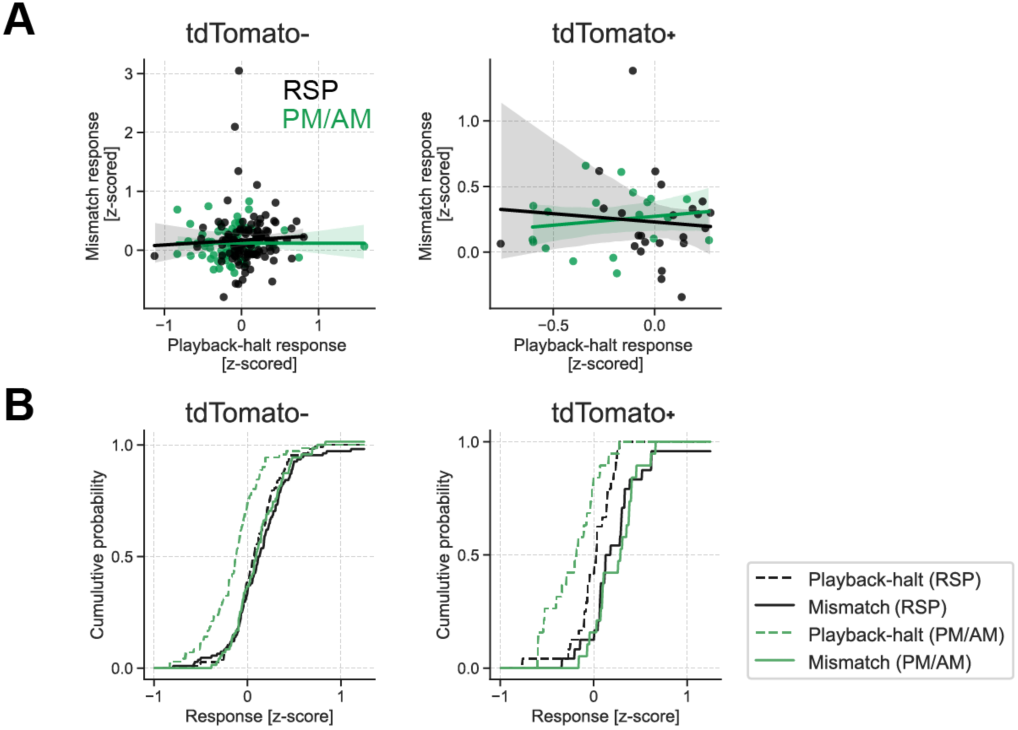
Mismatch responses of RSP and PM/AM neurons were not explained by visual flow halt responses, related to Figure 6. (A) Relationship between playback-halt responses and mismatch responses of individual neurons (a subset of neurons from Fig. 4). Solid lines and shaded regions represent linear regression lines and 95% CI. The magnitude of the playback-halt responses was not significantly correlated with mismatch response in any case (Pearson’s correlation coefficient: tdTomato^-^ RSP neurons, *p* = 0.625; tdTomato^-^ PM/AM neurons, *p* = 0.976; tdTomato^+^ RSP neurons, *p* = 0.702; tdTomato^+^ PM/AM neurons, *p* = 0.515). (B) Distributions of playback-halt and the mismatch response magnitudes.

**Figure S15.**
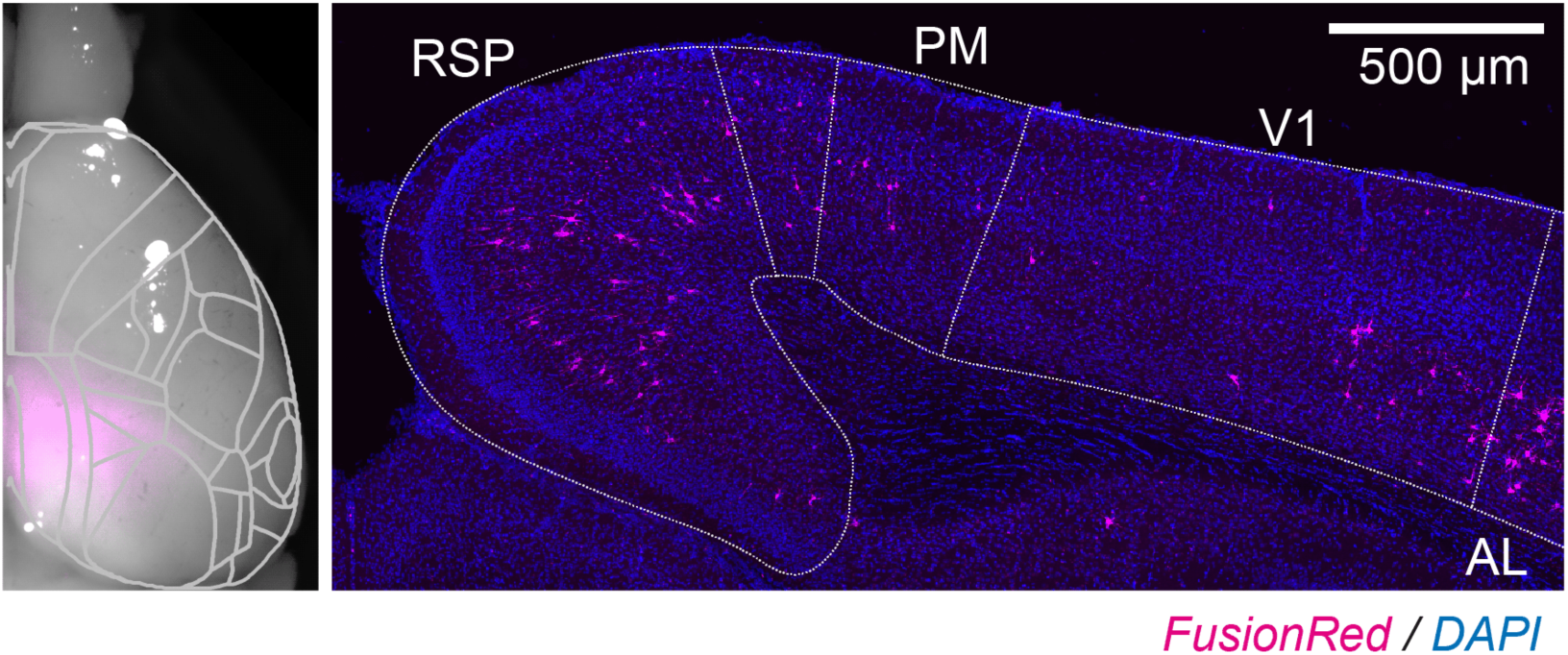
Cre-dependent expression of GtACR2 in the posterior cortex, related to Figure 6. Left: Fluorescence image of FusionRed (magenta) overlaid on a bright-field image (grayscale) with the ACCF; Right: Confocal image of FusionRed expression in the posterior medial cortex

**Figure S16.**
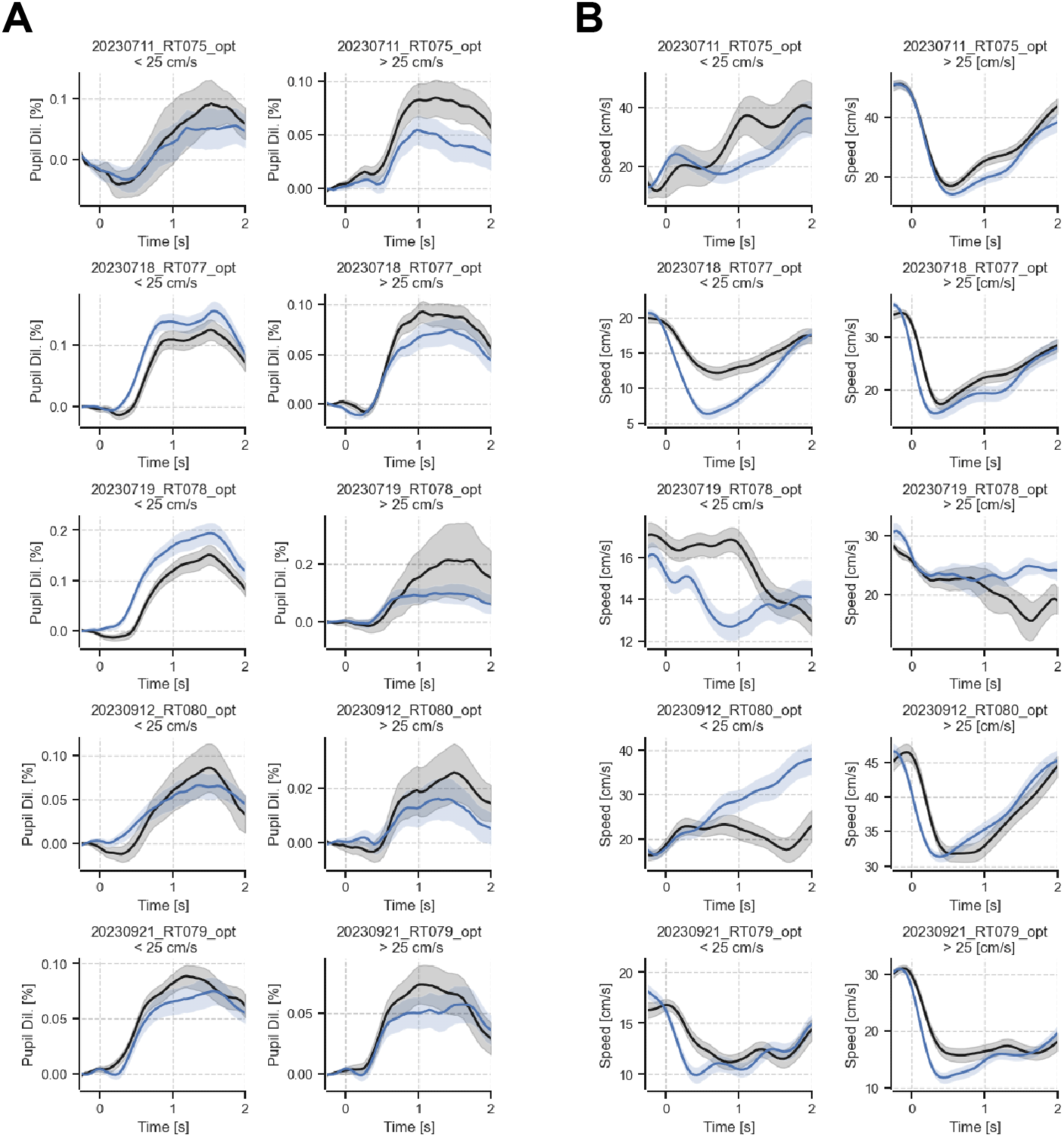
Effect of optogenetic manipulation in individual mice, related to Figure 6. (A). Effect of optogenetic silencing of ACC-projecting neurons in PM/AM and RSP on pupil responses in individual animals. Black lines indicate control trials. Blue lines indicate suppression trials. Shaded areas indicate SEM. (B). Effect of optogenetic silencing of ACC-projecting neurons in PM/AM and RSP on running speed in individual animals. Black lines indicate control trials. Blue lines indicate suppression trials. Shaded areas indicate SEM.

**Figure S17.**
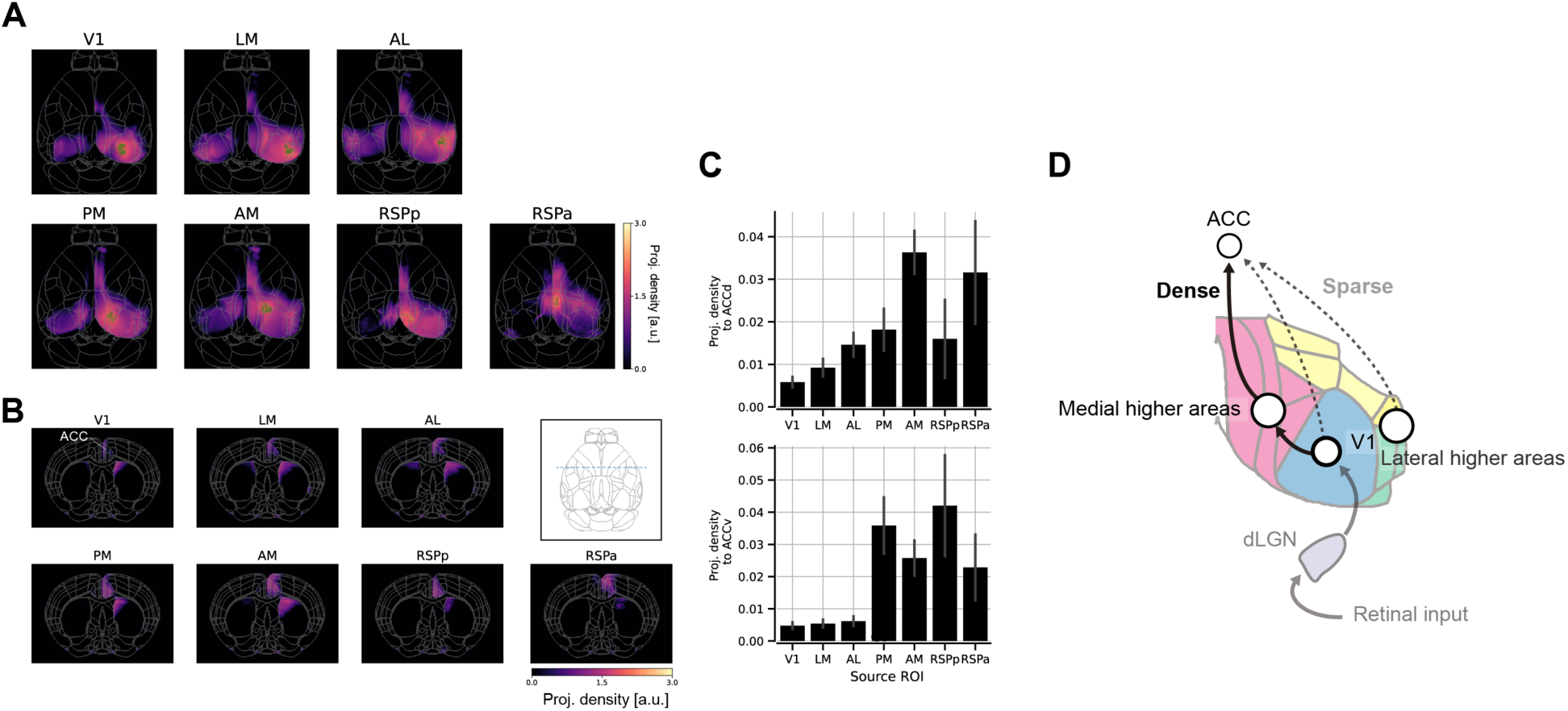
Axonal projections from posterior cortical areas to the ACC, related to Figure 6. (A). Top views of the ROI-seed axonal projection map in the Allen CCF. Green dots represent injection sites included in the analysis. (B). Coronal views of the ROI-seed axonal projection map. The inset shows the top view of the coronal coordinate (blue dashed line). (C). Axonal density in dorsal or ventral portions of ACC (ACCd or ACCv) for each experimental set. The error bar shows the SEM. (D). Summary of axonal projection from visual cortex and RSP to ACC.

**Figure S18.**
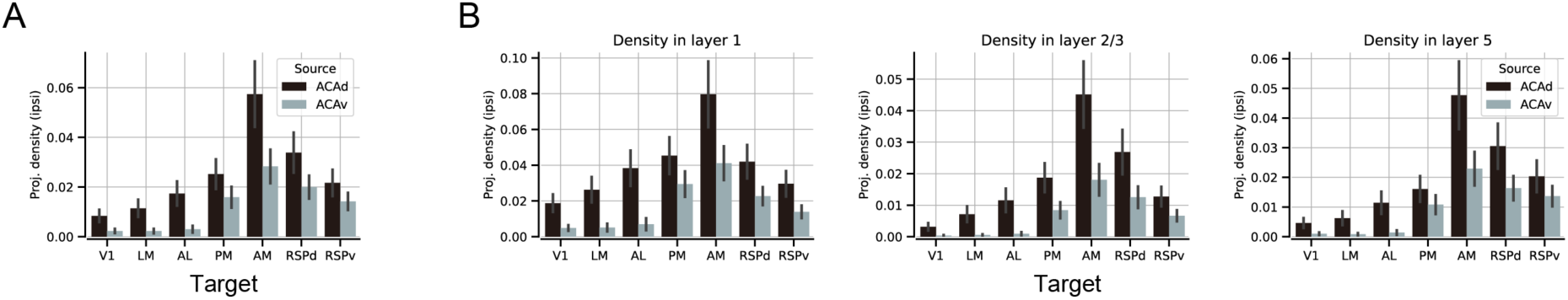
Axonal projections from ACC to posterior cortical areas, related to Figure 6. (A) Projection density from dorsal or ventral portions of ACC (ACCd/ACCv) to each cortical area. (B) Same as (A) but specifically for layer 1, layer 2/3, and layer 5 of each area.

**Supplementary Video 1. Resting-state activity across the cortex, related to Figure 1**

Wide-field imaging of a resting-state mouse. Top left: Timelapse of ΔF/F processed from raw fluorescence signals. Gaussian filtering was not performed. Top right: Face movie of an anesthetized mouse. Bottom left: Raw fluorescence movie acquired at 470-nm excitation. Bottom right: Raw fluorescence movie acquired at 405-nm excitation.

**Supplementary Video 2. Cortex-wide activity under closed-loop conditions, related to Figure 1**

Mouse running under the closed-loop condition. Brain activity, facial image, eye cropped image, pupil size, eye position, and running speed were displayed. Magenta arrows indicate the mismatch onset.

**Supplementary Video 3. Visuomotor mismatch responses across the cortex, related to Figure 1**

Trial-averaged movie of mismatch activities across cortical areas.

## Notes

### Competing Interest Statement

The authors have declared no competing interest.

### Summary of Updates

We have added electrophysiology, two-photon imaging, optogenetics, and modeling analyses in the revised manuscript.

